# Gibberellin transport affects (lateral) root growth through HY5 during Far-Red light enrichment

**DOI:** 10.1101/2023.04.21.537844

**Authors:** Kasper van Gelderen, Kyra van der Velde, Chia-kai Kang, Jessy Hollander, Orfeas Petropoulos, Tuğba Akyüz, Ronald Pierik

**Affiliations:** Plant-Environment Signaling, Dept. of Biology, Utrecht University, Padualaan 8 3584CH, Utrecht, The Netherlands; Light Signaling and Cell Biology, Centre for Organismal Studies, Heidelberg University, Im Neuenheimer Feld 230, 69120, Heidelberg, Germany

## Abstract

Plants compete for light by growing taller than their nearest competitors. This is part of the shade avoidance syndrome and is a response to an increase of Far-Red light (FR) reflected from neighboring leaves. The root responds to this shoot-sensed FR cue by reducing lateral root emergence. It is well-established that the plant hormone Gibberellic Acid (GA) is involved in supplemental FR-induced shoot elongation. Although GA is also transported from shoot to root, its role in regulating lateral root growth is unclear. We show via GA manipulations, both chemical and genetic, that GA modulates the lateral root reduction induced by shoot-sensed FR enrichment. Using the FRET-based GA biosensor GPS1, we observed detailed GA changes in the root upon shoot exposure to FR enrichment and when GA was supplied to the shoot. Supplying GA to the shoot also mitigated the FR-enrichment root phenotype, indicating a functional link between GA and changes in root development in response to shoot-sensed FR. The regulatory role of GA in root growth appears to be partially dependent upon the role of ELONGATED HYPOCOTYL 5 (HY5), a light-responsive transcription factor that regulates root growth. Shoot-to-root transported GA_4_ led to an increase in HY5 protein levels in the lateral root primordia. HY5 then repressed auxin signaling to repress lateral root growth. Our data unveil a novel way in which hormone and light signaling coordinate development across spatial scales by adjusting (lateral) root growth from above-ground FR light signals.

## Introduction

Light is an essential requirement for plant life and plants compete for light with other plants. Plants can detect competitors through reflection of far-red (FR) light from surrounding plants. This reflection increases the relative amount of FR, which is sensed through phytochromes (Franklin and Whitelam, 2005). Phytochromes can be activated by red (R) light and inactivated by far red light. In a high R:FR ratio, phytochrome B (phyB) represses the PHYTOCHROME INTERACTING FACTORS (PIFS), by binding and targeting them for 26S proteasome degradation (Shin et al., 2016; Ni et al., 2013). In a low R:FR ratio, indicative of neighbor proximity, phyB is inactivated and PIFs can activate shade avoidance responses of the shoot (elongation growth, early flowering (Ballaré and Pierik, 2017)). Low R:FR detected by the shoot can also lead to a reduction in lateral root density and main root length in Arabidopsis and this is regulated via the bZIP transcription factor ELONGATED HYPOCOTYL 5 (HY5) (van Gelderen et al., 2018).

Gibberellins (GA) are a class of growth promoting hormones that are also involved in the developmental responses of the shoot to Far-Red light (Bou-Torrent et al., 2014; Crocco et al., 2015; Djakovic-Petrovic et al., 2007; Hisamatsu et al., 2005). Low R:FR treatment enhances GA biosynthesis, while at the same time increasing the responsiveness of the plant to it (Hisamatsu et al., 2005; Reed et al., 1996). Gibberellin sensing occurs via the interaction between the GA sensor GIBBERELLIN INSENSITIVE DWARF1 (GID1) and DELLA repressor proteins (Schwechheimer, 2012). GA promotes GID1 binding to DELLAs which leads to the proteasomal degradation of the latter. DELLAs bind and repress the BRASSINAZOLE RESISTANT – AUXIN RESPONSE FACTOR – PIF (BAP) transcription factors module. These transcription factors together regulate cell expansion and thus also light-mediated cell elongation (Oh et al., 2014). The DELLA proteins form a brake on this BAP module, allowing GA-mediated degradation of DELLAs to release that brake and initiate shade avoidance responses (Daviere and Achard, 2016).

The shade avoidance response of the plant occurs throughout the entire plant body, but the sensing is confined to the leaves and stem of the plant, thus requiring a coordination between organs by signaling molecules (Küpers et al., 2018). Especially the response of the root system requires a long-distance communication between shoot and root, since the root does not sense the light when growing in dark soil. HY5 is involved in the root response to FR light detected by the shoot (van Gelderen et al., 2018) and can be transported over longer distances, affecting nitrate uptake and root growth (Chen et al., 2016; van Gelderen et al., 2021), phosphate and iron uptake (Sakuraba et al., 2018; Guo et al., 2021). In the root, HY5 acts by repressing auxin transport and signaling (Sibout et al., 2006; van Gelderen et al., 2018; Cluis et al., 2004), and it is specifically increased at the lateral root primordia when the shoot detects a low R:FR ratio (van Gelderen et al., 2018).

Gibberellins regulate shoot and root growth processes (Hedden and Sponsel, 2015). For the root, the concentration of GA needed to sustain growth is lower than the shoot (Tanimoto, 1994), and an overly high concentration can even slow down root growth (Inada and Shimmen, 2000). GA, through repressive effects on DELLAs, regulates the size of the root apical meristem (RAM) and root cell size, and GA acts in, and is synthesized in, the endodermis layer of the root tip (Barker et al., 2021; Ubeda-Tomás et al., 2009, 2008). Crucially, GA can also be transported from shoot to root (Binenbaum et al., 2018). Bioactive gibberellin, such as GA_3_ and GA_4_, can be traced from the shoot to the root tip through the phloem, where it unloads into the endodermis (Shani et al., 2013; Tal et al., 2016). Furthermore, it was shown that functional GA biosynthesis in the shoot is necessary to drive root growth and that even a precursor of bioactive GA, GA_12_, can be transported from shoot to root, where it is converted into bioactive GA (Regnault et al., 2015).

Here, we investigate how GA regulates root growth plasticity in response to aboveground FR cues of neighbor proximity. We show that shoot-to-root transport of GA can modulate the root response to supplemental shoot FR, and identify local GA changes using the GPS1 GA biosensor. Furthermore, we identify a role for HY5 in regulating the GA response and show that GA can affect HY5 abundance in the root.

## Results

### GA and paclobutrazol application modify effect of supplemental FR on lateral root development

In order to investigate whether GA plays a role in the response of the root system to supplemental Far-Red light we performed dose-response experiments. We illuminated the shoot with supplemental Far-Red light (WL+FR) to lower the R:FR ratio and used D-root plates (Silva-Navas et al., 2015) to avoid direct illumination of the root system (van Gelderen et al., 2021, 2018). Seeds were germinated on ½ MS control media, transferred after one day to either control or GA-containing ½ MS media (Supplemental Figure 1A), and transferred after four days to new plates (van Gelderen et al., 2018). We used the GA_4_ form of Gibberellic acid, since it is a more prevalent bioactive form in Arabidopsis than the often used GA_3_ (Prasetyaningrum et al., 2020), with a good affinity to the gibberellin sensor GID1 (Yoshida et al., 2018). However, we compared its effect against that of GA_3_, which confirmed that GA_4_ is a more potent gibberellin than GA_3_ (Supplemental Fig. 1). Thus, we continued the use of GA_4_ in subsequent experiments. GA_4_ application led to increased hypocotyl growth and an increased response to WL+FR, with a maximum response at 10^2^ nM of GA_4_ (Fig. 1a). Up to 10^2^ nM GA_4_, main root growth was unaffected, however at 10^3^ nM there was an increase in main root length (Fig. 1d). Interestingly, 10^4^ nM of GA_4_ led to a strong decrease in main root length, coupled to an increase in root hair formation, and agravitropic root growth (Fig. 1d,e). Lateral root density in control treatment decreased due to WL+FR, but GA_4_ application at 10^0^-10^2^ nM negated this effect (Fig. 1b,e) (except at 10^3^ nM GA_4_) of GA_4_, while at 10^4^ nM very few lateral roots emerged at all (Fig. 1d,e). WL+FR decreased the lateral root zone length,also called the branching zone, which is the length of the main root between the first and last emerged initial (Fig. 1c). GA_4_ application removed the negative effect of WL+FR on lateral root zone length, varying with the concentration of GA_4_ (Fig. 1c).

**Figure 1.**
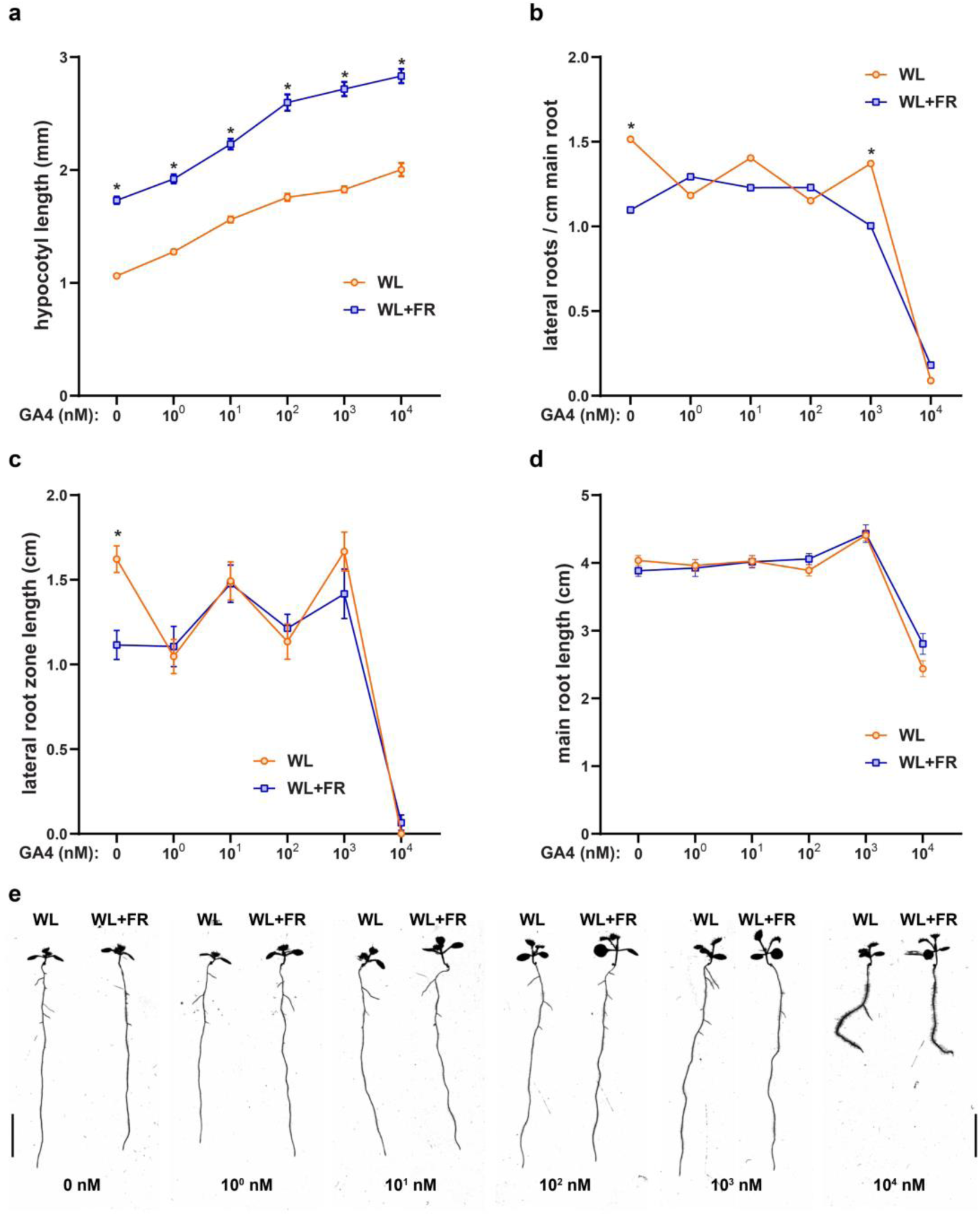
GA_4_ application modifies the response of the root to supplemental Far-Red light. Seedlings were grown for 9 days according to the schedule on Fig. S1a. Scans of 9d old seedlings were analyzed on (a) hypocotyl length; (b) main root length; (c) lateral root density; (d) lateral root zone length. (e) Representative seedling images of the experiment, scale bar = 1 cm. Means were statistically significant based on a 2-way ANOVA; letters denote significant difference between treatments based on a post-hoc tukey test (p<0.05).

Paclobutrazol (PAC) is a drug that inhibits the conversion of *ent-*Kaurene into *ent-*Kaurenoic acid, a crucial first step in gibberellin biosynthesis that is performed by the GA1 enzyme (Rademacher, 2000). Genetically knocking out GA1 leads to a dramatic loss of GA production, however by dosing the amount of PAC one can gradually inhibit biosynthesis of GA. We grew Col-0 seedlings in a dose-response experiment similar to the one presented in Fig. 1, however now we used 0, 10^2^, 10^3^ and 10^4^ nM of PAC. PAC treatment led to a decrease in WL+FR-induced hypocotyl elongation that was completely removed at 10^4^ nM PAC (Fig. 2a). 10^2^ nM PAC reduced the lateral root density and prevented the reduction induced by WL+FR (Fig. 2b). Lateral root density was further reduced by 10^3^ nM PAC, however, at 10^4^ nM it rose sharply (Fig.2b); This strong increase can be attributed to a dramatic decrease in main root length (Fig. 2c,d), thus concentrating the lateral roots on a shorter main root. Interestingly, while the main root was smaller than 1 cm on average in this very high PAC concentration, still some lateral roots could emerge (Fig. 2d). Additionally, we tested the *ga1-3* mutant, which is defective in the GA1 biosynthesis enzyme, and found that the WL+FR response was lost in this mutant (Fig. S1f). Both the addition of GA_4_ and PAC to the growth medium leads to the loss of the effects brought on by the WL+FR treatment. Therefore, we conclude that gibberellins can modulate the shoot and root response to shoot-applied FR light.

**Figure 2.**
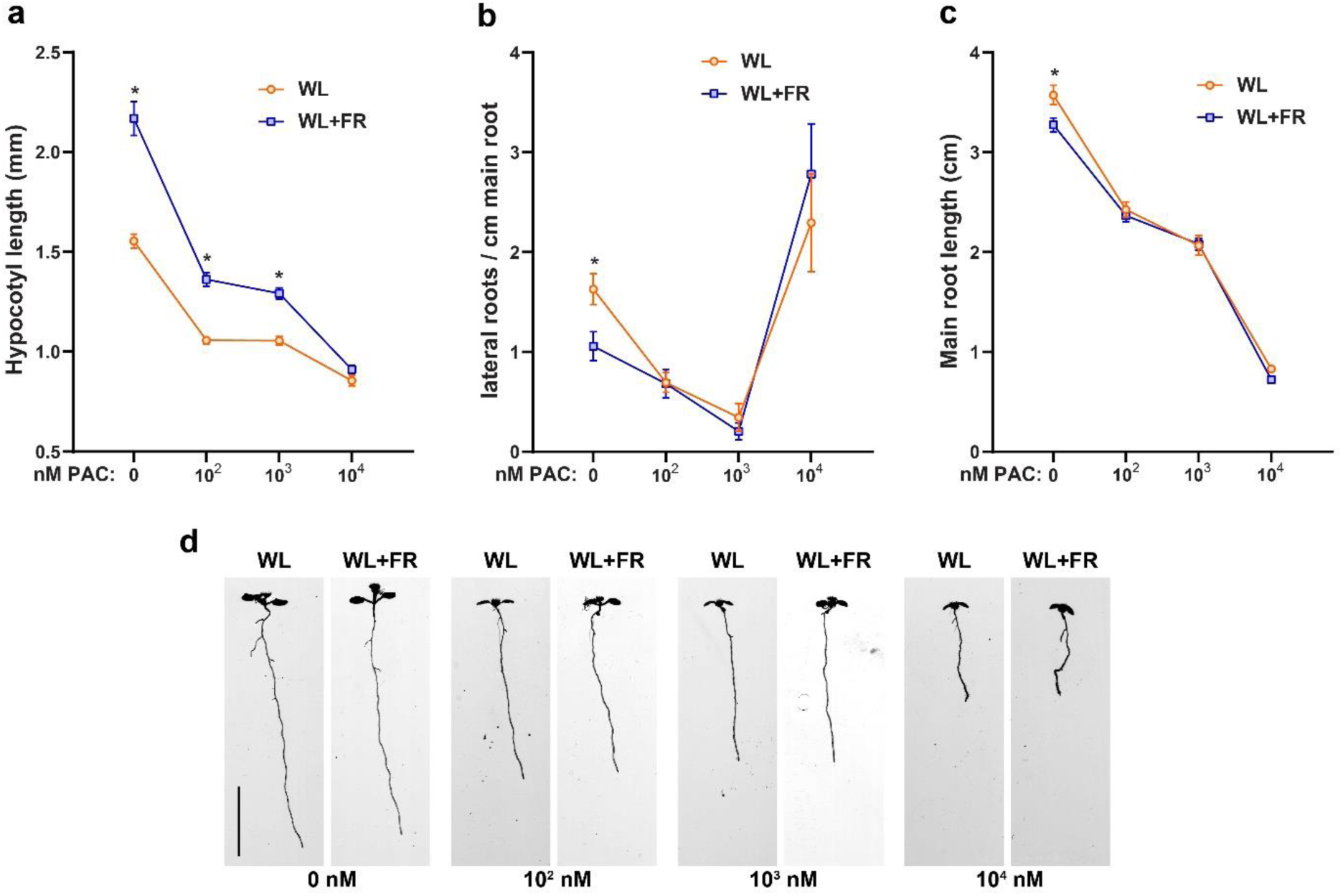
Paclobutrazol treatment reduces root growth and removes the effects of WL+FR on hypocotyl and root development. Seedlings were grown for 9 days according to the schedule on Fig. S1a. Scans of 9d old seedlings were analyzed on (a) hypocotyl length; (b) lateral root density; (c) main root length; (d) Representative seedling images of the experiment, scale bar = 1 cm. Means were statistically significant based on a 2-way ANOVA; letters denote significant difference between treatments based on a post-hoc tukey test (p<0.05).

### Supplemental FR light to the shoot increases GA in the root

Since GA_4_ addition leads to the loss of WL+FR effects on lateral root development, we chose to investigate whether WL+FR would lead to changes in the levels of GA in the root system. To do this we used the GPS1 Förster Resonance Energy Transfer (FRET) sensor, that is able to detect relative levels of active Gibberellins by measuring the ratio in YFP to CFP signal that are fused to a truncated GID1 and DELLA (GAI) (Rizza et al., 2021, 2017)(Fig. 3a). Four-day old seedlings were transferred to control or 10^2^ nM GA_4_ plates and after two days were fixed via the clearsee method (Ursache et al., 2018). First, we checked if the GPS1 sensor was responsive in the root to GA_4_ addition to the plate medium. Both in the Root Apical Meristem (RAM) and the lateral root primordium did we see a pronounced increase in the YFP/CFP ratio when 10^2^ nM of GA_4_ was applied (Fig. 3b), which corresponds to an increase in the bioactive GA level. The non-responsive control sensor GPS1-NR did not react to external GA application (Fig. 3c). A treatment of WL+FR during two days led to an increase in the GPS1 ratio in the elongation zone of the root meristem (Fig. 3d,g-h), which is interesting since the elongation zone has been identified as a region where shoot-borne GA_4_ can be transported to (Shani et al., 2013). In the lateral root primordia we also observed an increase in the YFP/CFP GPS1 ratio, across all stages (Fig. 3e), and mainly in stages 4 and 5 (Fig. 3f,i-j). These GPS1 data show that WL+FR, as detected by the shoot, leads to an increase in GA levels in root tissues, particularly around the elongation zone and in the lateral root primordia, which are regions important for main root length and lateral root density respectively.

**Figure 3.**
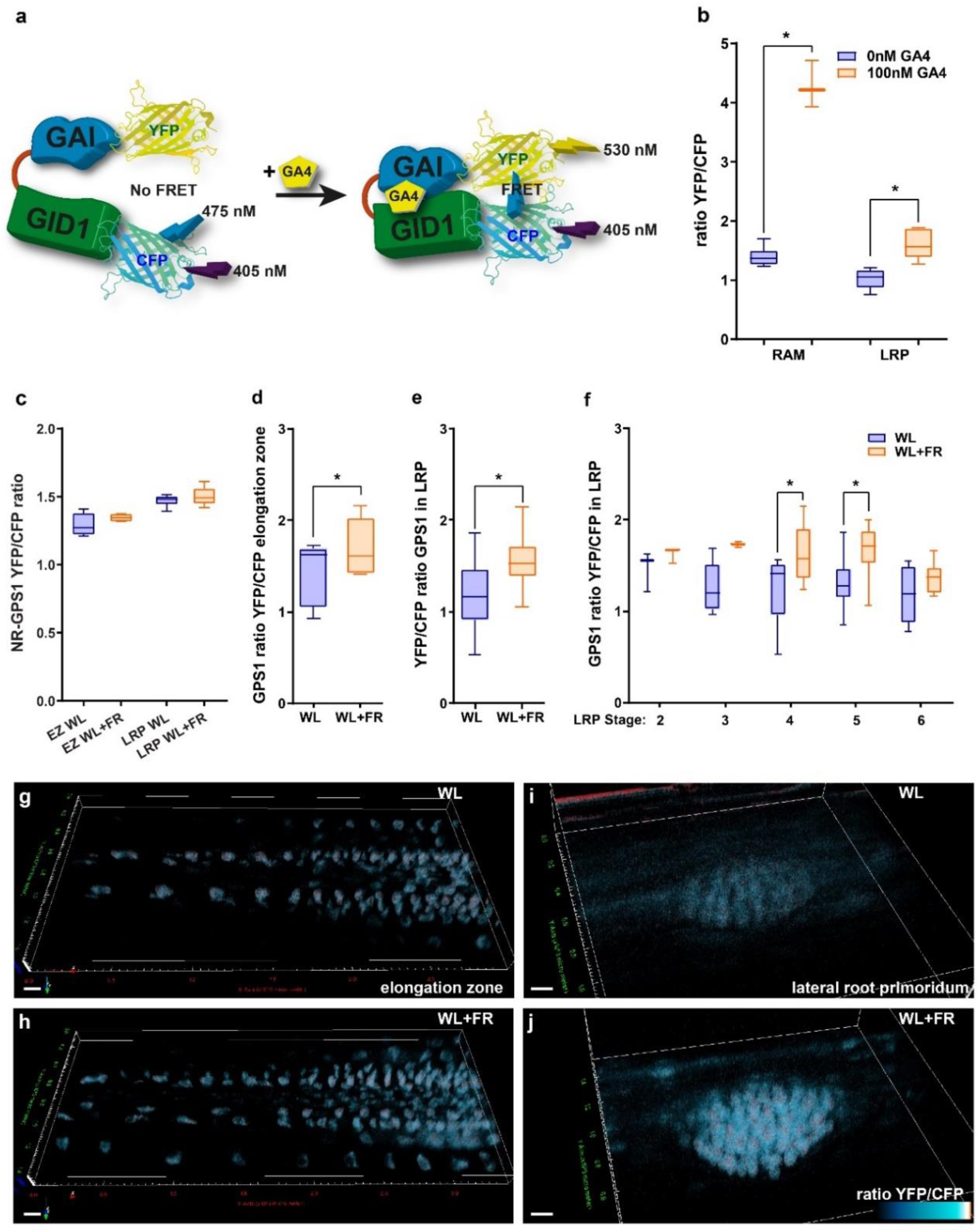
WL+FR leads to increased GA as detected via the GPS1 sensor. Confocal microscopy experiment with quantifications (b-f) and representative images (g-j). (a) Cartoon explaining the principle behind detecting bioactive GA with GPS1 (based on Rizza et al., 2017). (b) Ratio between YFP and CFP emission of GPS1 in six day old seedlings treated with mock or GA_4_ for two days and similar data for the non-responsive (NR) variant of GPS1 (c). (d-f) GPS1 ratio in 6 day old seedlings treated with WL or WL+FR for five days; *significant difference based on student’s t-test (d,e) and one-way ANOVA (f) (p<0.05). (g-j) representative YFP/CFP ratiometric 3D-rendered images of experiments in (d-f); scale bar = 10 μm.

### GA4 application to the shoot reduces root growth and removes WL+FR effect on root development

Several reports have alluded to potential roles for GA transport from the shoot to the root in affecting root development (Binenbaum et al., 2018; Shani et al., 2013) (Regnault et al., 2015). We detected an increase in GA in WL+FR via the GPS1 sensor. Since in our setup, WL+FR is only detected by the shoot, these findings could suggest that WL+FR leads to increased transport of GA from shoot to root. In order to test this notion, we wanted to see if shoot-applied GA_4_ can lead to root developmental changes. Therefore, we designed square petri-plates where we could separate a shoot and root compartment by a physical barrier of 3mm height, so that different layers of agar could be poured but not diffuse into each other (Fig. S2).

We grew seedlings for four days on normal square plates on ½ MS medium without GA_4_ and then transferred them to the two-compartment plates containing control (EtOH 1:10000) or 10^2^ nM GA_4_ medium in the respective compartments (Fig. 4a). We took extra care to position the seedlings such that the hypocotyl and cotyledons, but not the root touched the medium in the upper compartment and we carefully placed the root over the plastic barrier, so that it touched the bottom compartment (Fig. S2c). We used a full factorial combination of GA and control (EtOH) in the top or bottom compartment (EtOH/EtOH, GA_4_/EtOH, EtOH/GA_4_, GA_4_/GA_4_). After 5 days of growth on the compartment plates we scanned and analyzed hypocotyl length and root traits (Fig. 4b). Hypocotyl length was increased only on the plates which contained GA_4_ medium in the top compartment (GA_4_/EtOH and GA_4_/GA_4_) (Fig. 4c).

**Figure 4.**
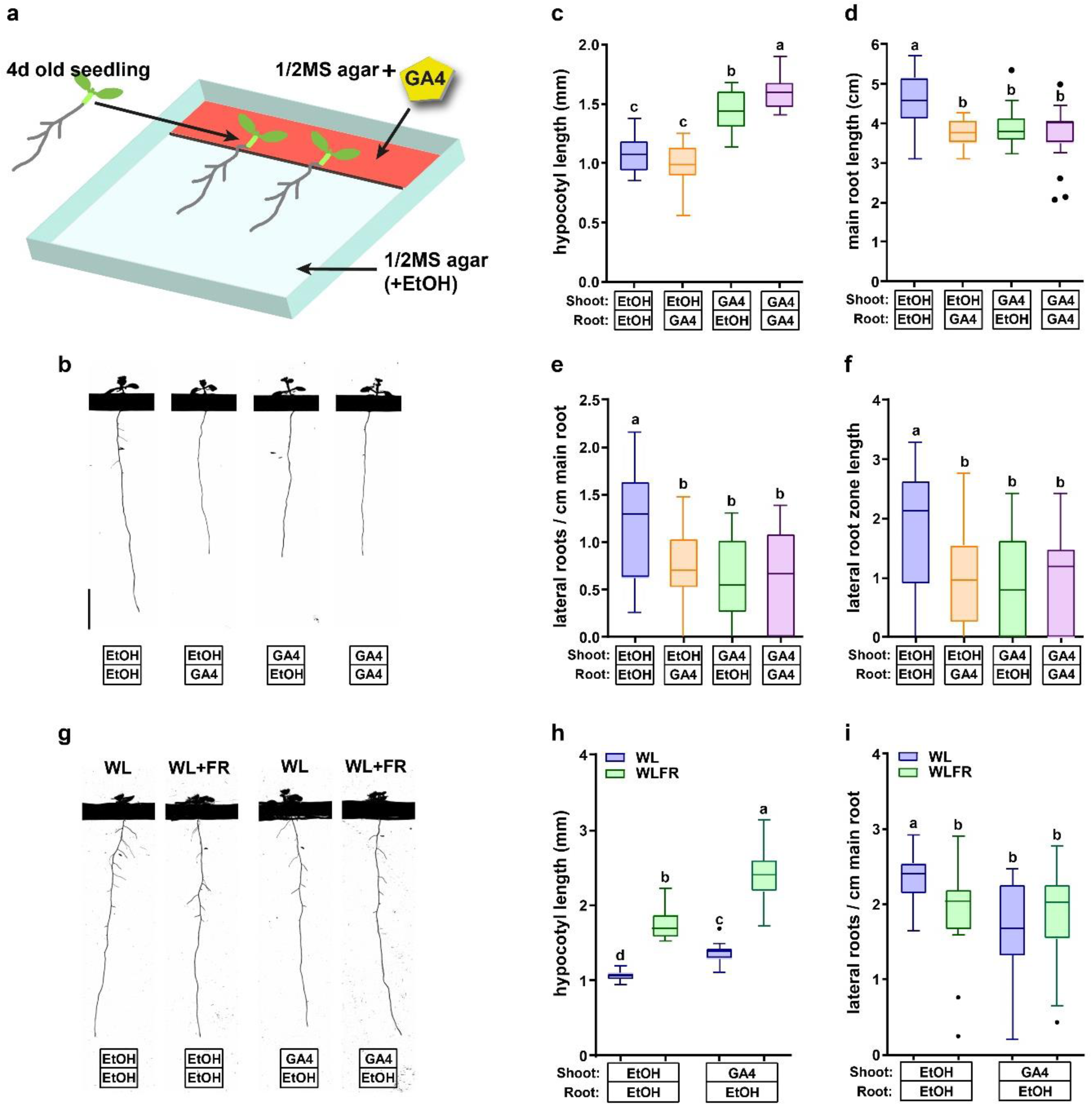
Compartmentalized plates show that shoot applied 10^2^ nM GA_4_ affects (lateral) root growth similarly to 10^2^ nM GA_4_ applied directly at the root. (a) Overview of the experimental procedure. A barrier of 3mm high is present on the plate that physically isolates two agar layers. The top layer is 3cm long, while the bottom one is 9cm. Seedlings are first grown on normal ½ MS for four days, and then they are transferred so that the start of root is carefully positioned over the barrier. (b-f) Nine day old Col-0 wild type seedlings grown in WL on compartmentalized plates with different media combinations. (b) representative seedlings, black bar is the physical barrier on the plate, scale bar = 1cm; (c) hypocotyl length, (d) main root length; (e) lateral root density; (f) lateral root zone length. (g-i) Ten day old Col-0 wild type seedlings grown in WL or WL+FR on medium with or without GA_4_ in the top compartment. (g) representative seedlings, scale bar = 1 cm; (h) hypocotyl length; (i) lateral root density. Means were statistically significant based on a 2-way ANOVA; letters denote significant difference between treatments based on a post-hoc tukey test (p<0.05).

However, main root length was negatively affected by GA_4_, irrespective of it being in the top or bottom compartment or both (Fig. 4d). Similarly, lateral root density and lateral root zone length were negatively affected in the same manner if GA_4_ was given directly to the bottom compartment or only to the shoot (GA_4_/EtOH, EtOH/GA_4_) and GA_4_/GA_4_) (Fig. 4e-f). This shows that supplying the root with GA_4_ does not lead to hypocotyl elongation, however, when the shoot is in contact with GA_4_ medium the effect on the root system is the same as when the root itself is in GA_4_ medium. This indicates that shoot-to-root transport of GA_4_ may play a role in changing main and lateral root development, although an intermediate mobile messenger cannot be ruled out yet.

Next, we tested whether application of GA_4_ to the top compartment would remove the effect of WL+FR on lateral root growth, similar to application on the whole plate (Fig.1b). After the 4d old seedlings were transferred to either an EtOH/EtOH or a GA_4_/EtOH plate, these plates received either WL or WL+FR (Fig. 4g). Hypocotyl length after six additional days of growth was increased on GA_4_/EtOH plates and also by WL+FR, in a manner very similar to GA_4_ application on the whole plate (Fig. 4h and 1a).

WL+FR led to a reduction in lateral root density on EtOH/EtOH plates, as did GA_4_/EtOH, WL+FR did not affect lateral root density any more on GA_4_/EtOH plates (Fig. 4i). In all these experiments of Figure 4, 10^2^ nM GA_4_ application led to a reduction in main and lateral root growth, contrary to the results in Fig.1. This is possibly due to the different timing of GA_4_ exposure: four days after germination in Fig. 4, versus one day after germination in Fig. 1. Overall, the results on these compartmentalized plates indicate that shoot-borne GA_4_ can affect lateral root development and can modulate the response to shoot perceived WL+FR. Next, we wanted to know how this application of GA_4_ on the shoot would affect the levels of GA as measured by GPS1.

### GA4 applied on shoot leads to significant changes in bioactive GA in the root

In order to assess the relative GA levels, we performed a similar experiment as shown in Fig. 4, however this time we used GPS1 seedlings that were fixed and cleared with clearsee 2d after transfer to preserve them for confocal microscopy imaging (Fig. 5). We observed that GA_4_ application to the shoot compartment led to an increase in in the YFP/CFP GPS1 ratio in the hypocotyl (Fig. 5a,e-f). Then, moving down towards the root tip we observed a significant increase in GPS1 ratio in the lateral root primordium due to GA_4_ application to the shoot (Fig. 5b,g-h). Importantly the GPS1 ratio was only increased in the inner layers of the root that are enclosed by the casparian strip of the endodermis: The stele, pericycle, and endodermis, while the cortex and epidermis layers did not show an increase of the GPS1 ratio (Fig. 5g-j). In comparison, when GA_4_ was applied to the whole plate the GPS1 ratio ncreased also in the cortex and epidermis (Fig. S3). Again, in the maturation zone the GPS1 ratio increased strongly in the stele+pericycle+endodermis, and not the cortex and epidermis (Fig. 5c,i-j). This increase only in the tissue enclosed by the casparian strip would be consistent with GA being transported from shoot to root, through the vascular tissue. In the elongation zone, where the Casparian strip is not yet developed (Barberon, 2017), and the phloem unloads into the root tissue (Ross-Elliott et al., 2017), we observed an increase in GPS1 ratio across all layers of tissue (Fig. 5d,i-j). This is in accordance with the observed shoot-root transport of fluorescent GA_3_ and GA_4_ (Shani et al., 2013).

**Figure 5.**
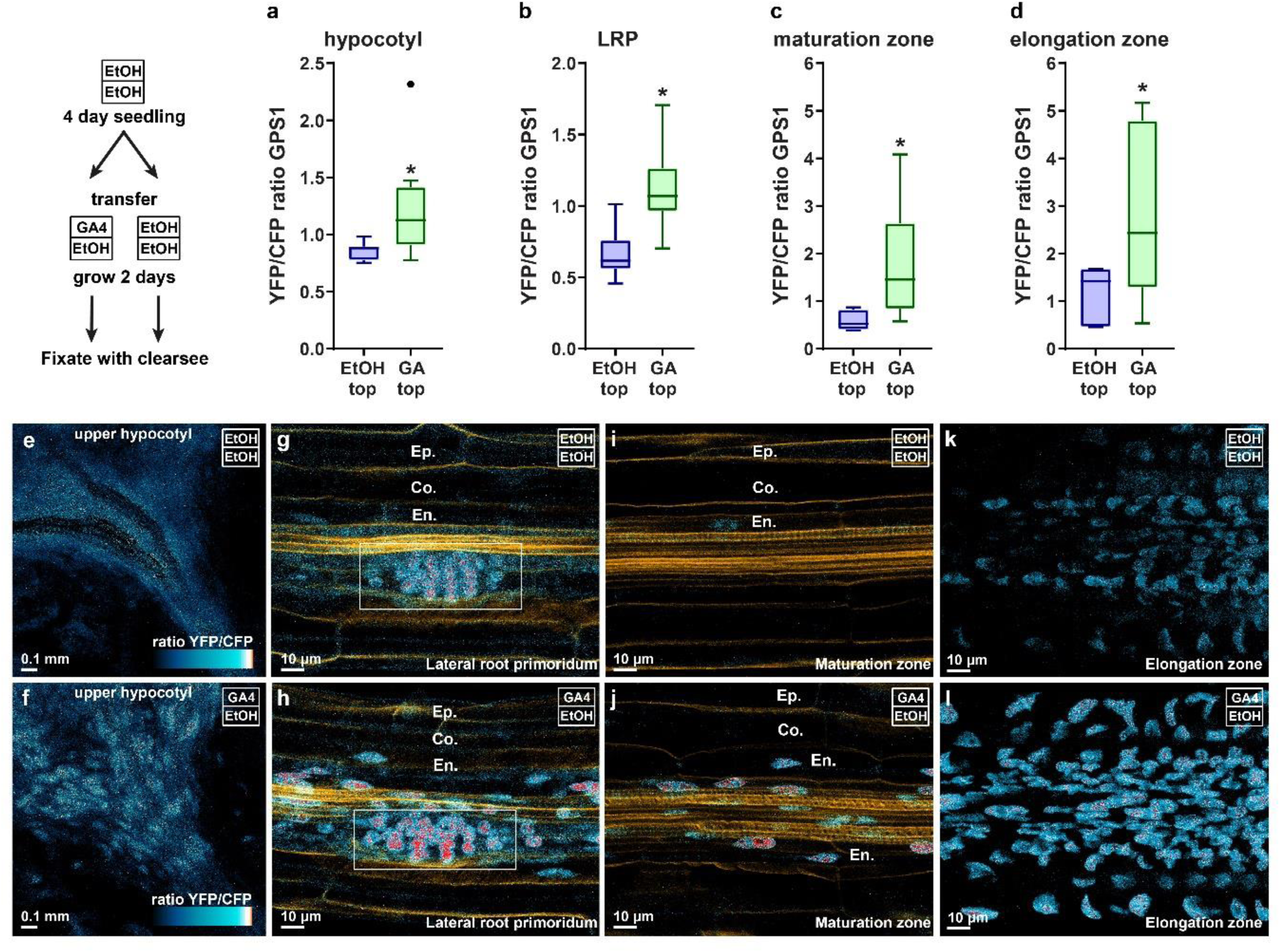
Shoot application of GA_4_ leads to strictly defined GPS1 FRET changes in the root. Confocal microscopy experiment with quantifications (a-d) and representative images (e-l). GPS1 seedlings were handled in the same manner as depicted in Fig. 4a, with the difference being that the seedlings were fixed with clearsee after two days. GPS1 line was either treated for two days with shoot GA4, or with mock EtOH. (a-d) GPS1 YFP/CFP ratios of the hypocotyl (a), lateral root primordia (b), maturation zone (c), elongation zone (d). (e-i) Representative images used for calculating (b): (e-f), (c): (g-h), (d): (i-j). White box emphasizes the LRP, and the epidermis (Ep.), cortex (Co.) and endodermis (En.) layers have been highlighted in g-i and a cell-wall stain with directred23 is visible in orange. Scale bars = 5 μm. *significant difference based on student’s t-test (p<0.05).

Summarizing, we show that WL+FR treatment of seedling shoots leads to increased GA levels in the lateral root primordia and the main root elongation zone, and that GA_4_ application reduces lateral root density and lateral root zone length similar to WL+FR. Finally, we confirm that shoot-applied GA is likely transported to the root, which indicates that GA could play an important role in transducing the WL+FR signal from the shoot to the root. Next, we investigated the downstream regulation following GA_4_ application and how this integrates with the regulation of lateral root growth by WL+FR.

### DELLAs confer the GA effects on the root in WL+FR

In order to understand how GA is involved in the regulation of lateral root development by WL+FR we started with the knockout mutant of the five *della* GA repressor proteins, which should have very little response to GA_4_ application since GA responses typically occur via GA-dependent DELLA protein degradation (Schwechheimer, 2012). Compared to the L*er* background, the *della pentuple* mutant had an increased hypocotyl length in WL and WL+FR (Fig. S4a, consistent with a previous study (Hayes et al., 2019)), but did not respond further to GA application, as expected (Fig. S4a,d). L*er* showed a reduction in lateral root density in WL+FR and in accordance with Col-0 this reduction was no longer present in the 10^2^ nM GA_4_ treatment (Fig S4b,d). Importantly, the lateral root density and lateral root zone length of the *della pentuple* mutant was not affected by WL+FR, or GA_4_ treatment (Fig. S4b-d). We previously showed that the lateral root primordium stage 5 and 6 are enriched in WL+FR, consistent with a reduced lateral root emergence (van Gelderen et al., 2021, 2018) and this phenotype was also lost in the *della pentuple* mutant (Fig. S4e). Since DELLAs are degraded upon binding of GA to its receptor GID1 (Schwechheimer, 2012), we tested whether our 10^2^ nM GA_4_ treatment effectively degraded DELLAs in the root. Protein levels of the DELLA protein RGA fused to GFP, visualized through confocal microscopy, were clearly visible in mock treatment, and disappeared after three days of 10^2^ nM GA_4_ (Fig. S5a), confirming that this GA_4_ treatment was sufficient to degrade one of the major DELLAs. We subsequently used an anti-RGA antibody to observe the RGA response to WL+FR. In accordance with our expectations and data, WL+FR led to a decrease in RGA in the shoot and importantly also the root (Fig. S5b,d) and as expected, 10^3^ nM 24h PAC treatment stabilized RGA (Fig. S5c). These results show that the DELLAs are important for the root response to shoot-exposed WL+FR and that they confer the effect of GA on the root system in WL+FR.

### Interactions between GA and HY5 regulate root responses to WL+FR

A previously established regulator root developmental response to WL+FR is the HY5 transcription factor (van Gelderen et al., 2018). We, therefore, investigated if the HY5 and GA pathways jointly regulate root development in response to light, using the *hy5 hyh* (*hy5 homolog*) double mutant. WL+FR and GA_4_ application both increased the hypocotyl length of Col-0 as well as *hy5 hyh* double mutant (Fig. 6a). The interactive effects of WL+FR and GA_4_ application on root development observed in Col-0 were absent in the *hy5 hyh* mutant (Fig. 6c-d). Since GA_4_ effects are particularly strong at high concentrations, we also performed an experiment with 10^4^ nM GA_4_, instead of 10^2^ nM. The strongly increased hypocotyl length, inhibition of main root length, reduced lateral root density and lateral root zone length and the increase in root hairs found in Col-0 (Fig. 6h-k), were partially absent in *hy5 hyh,* indicating a substantially reduced sensitivity of this mutant to GA application. *hy5 hyh* lateral root density was decreased by 10^4^ nM GA_4_, however, this was mainly due to the extra main root length, as the lateral root zone length strikingly did not change (Fig. 6i,j,l). Overall, the roots of *hy5 hyh* seedlings looked very similar in 10^4^ nM GA_4_ compared to 0 nM GA_4_ (Fig. 6l), in stark contrast to Col-0, which had almost no laterals left and many, very long root hairs (Fig. 6k,l, see arrowhead). We used the RGA antibody to see if this lack of GA sensitivity in *hy5 hyh* would be visible at the protein level. The decrease in RGA detected in WL+FR in Col-0, was not visible in the *hy5 hyh* double mutant (Fig. S5d), confirming that the *hy5 hyh* mutant is indeed less sensitive to GA.

**Figure 6.**
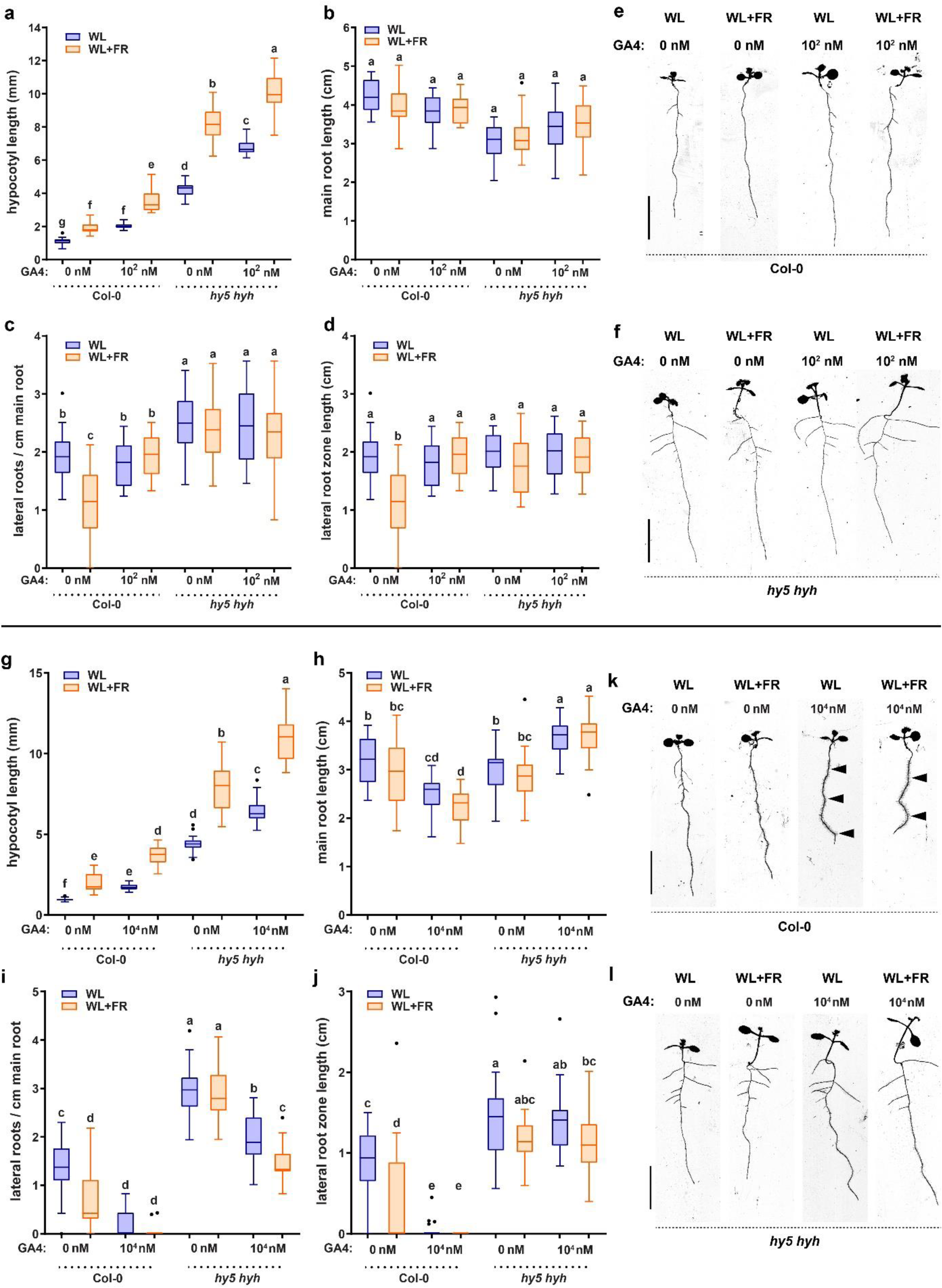
The *hy5 hyh* mutant has no lateral root development reduction in WL+FR nor in GA_4_ treatment and responds less severely to a very high dose of GA4. Seedlings of Col-0 and *hy5 hyh* were grown for 9 days according to the schedule on Fig. S1a. Scans of 9d old seedlings treated with EtOH 1:10000 in the medium or 10^2^ nM GA_4_ and analyzed on (a) hypocotyl length; (b) main root length; (c) lateral root density; (d) lateral root zone length. (e,f) Representative seedling images of the experiment in (a-d), the arrowheads in (k) point to excess root hairs in the 10^4^ nM GA_4_ treatment. (g-l) Similar setup as in (a-f), but then with 10^4^ nM GA4. scale bar = 1 cm. Means were statistically significant based on a 2-way ANOVA; letters denote significant difference between treatments based on a post-hoc tukey test (p<0.05).

Next, we also investigated the reverse interaction: effects of GA application on HY5 abundance, using a *hy5-2 pHY5:HY5-GFP* line. We treated seedlings for two days with 10^2^ nM GA_4_ and EtOH mock control, after which they were analyzed on the confocal microscope. Across all lateral root primordia stages there was a significant increase in HY5-GFP abundance in response to GA_4_ application (Fig. 7a,b). Conversely, inhibiting endogenous GA levels with paclobutrazol (PAC) did not significantly affect HY5-GFP abundance (Fig. 7c,d). We then treated seedlings with GA_4_ on the shoot and analyzed HY5-GFP in the shoot and lateral root primordia. GA_4_ shoot treatment caused a similar increase of HY5-GFP in the lateral root primordia as did the whole-plate GA_4_ treatment (Fig. 7b,e,f). Interestingly, HY5-GFP in the shoot apical meristem was decreased by GA_4_ application (Fig. 7g,h).

**Figure 7.**
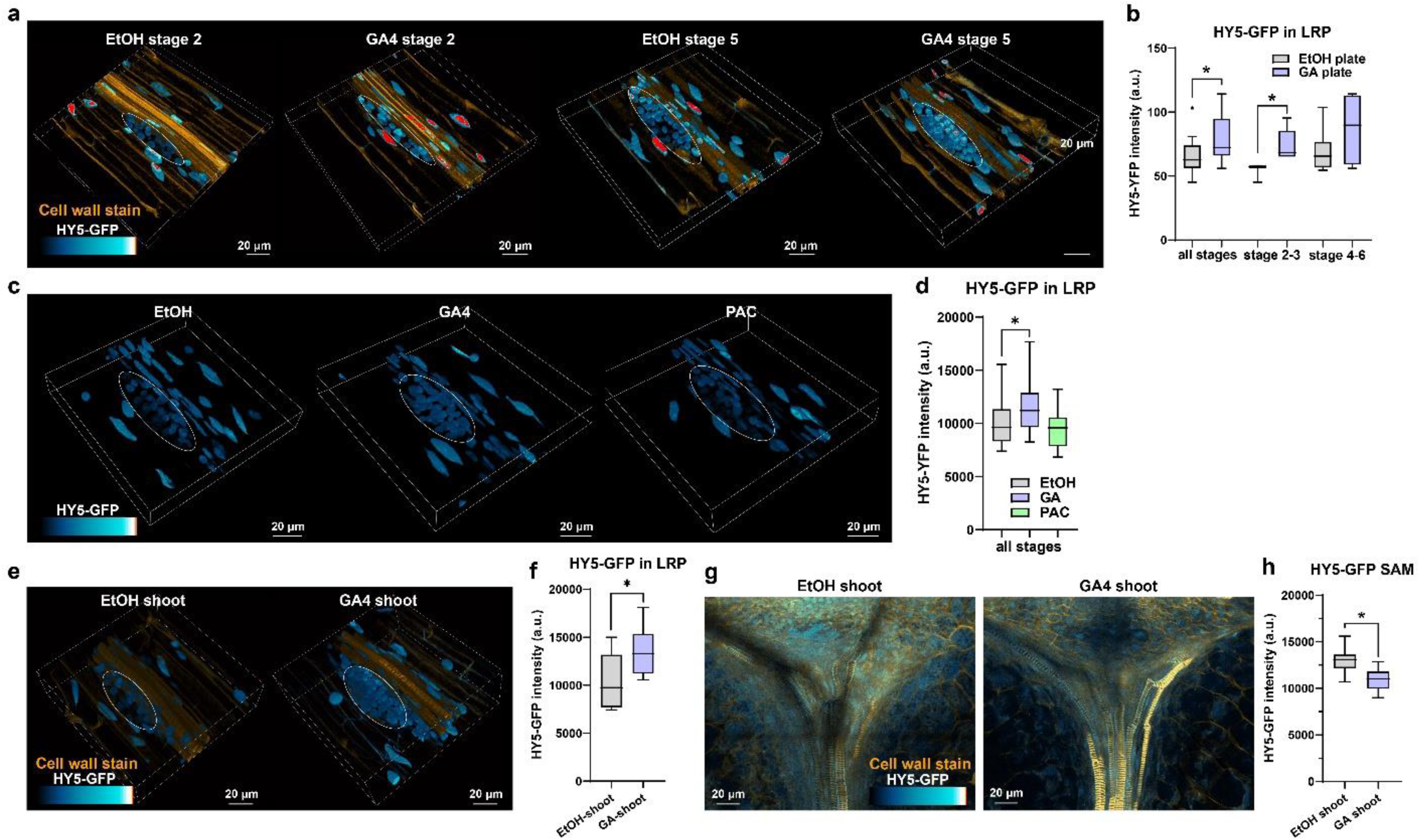
GA application leads to an increase in HY5-GFP in the root. Six day old *hy5-2 pHY5:HY5-GFP* seedlings treated with mock or GA_4_ on the whole plate for two days (a-d), or on the shoot for two days, similar to Figures 4 and 5 (e-h). (a,c,e) 3D projections of representative lateral root primordia with HY5-GFP in blue and, for (a) and (e), a direct red23 cell wall staining in orange. (b,d,f,h) Accompanying quantifications to (a),(c),(e) and (g) respectively. (g) 2D representative max intensity projections of the shoot apical meristem (SAM) region with HY5-GFP in blue and a direct red23 cell wall staining in orange. *significant difference based on student’s t-test (f,h) and one-way ANOVA (b,d) (p<0.05).

This striking difference between shoot and root HY5-GFP responses to GA_4_ application is consistent with the opposite effects of GA_4_ on growth in the two different tissues: GA_4_-induced reduction of HY5 would promote hypocotyl elongation, since HY5 represses hypocotyl growth in shade (Ortiz-Alcaide et al., 2019; Sellaro et al., 2011)(Fig. 4c), while the GA_4_-induced promotion of HY5-GFP in the root would lead to a shorter, reduced root system (van Gelderen et al., 2018) (Fig. 4d). HY5 can repress auxin signaling (Cluis et al., 2004; van Gelderen et al., 2018), and GA can post-translationally upregulate auxin transporters (Willige et al., 2011; Mäkilä et al., 2023). Therefore, we used the C3PO auxin sensor cassette (Küpers et al., 2023) to estimate auxin abundance and auxin response. In the lateral root primordia of stages 1-4, both auxin abundance and response, as measured by the R2D2 and DR5v2 parts of the sensor respectively, were promoted by shoot-GA treatment (Fig. S6a-c). In the phloem unloading zone, auxin levels increased due to shoot GA (Fig. S6d,e), while in the main root meristem, auxin signaling was reduced by shoot-GA treatment (Fig. S6f,g). Overall, GA_4_-shoot treatment led to an increase in auxin and though these data show that GA can crosstalk with auxin signaling, they also highlight the complexity of interactions between GA_4_, HY5 and auxin and future studies are needed to resolve if and how auxin regulation by GA and HY5 either or not is functionally associated with root responses to shoot-exposed FR light.

### HY5 and shoot-GA_4_ repress auxin signaling and lateral root development-related genes

To obtain insight into the downstream transcriptional effects of shoot-GA_4_ treatment and the role of HY5 in this process we performed a qPCR analysis on shoot and root samples of seedlings treated with shoot-GA_4_ or control ethanol (1:10000) in either wild type or *hy5 hyh* mutant background. We tested genes known to be downstream of or associated with HY5 (*NRT2;1, BBX21* (Chen et al., 2016; Datta et al., 2007; van Gelderen et al., 2021; Xu et al., 2016)), GA biosynthesis and signaling genes (*GA3ox1*, *GA2ox6*, *GA2ox8* (Hedden and Thomas, 2012) and RGA), auxin transport and signaling genes (*PIN3, LAX3, ARF19, SHY2, IAA2* (Li et al., 2015; Vilches-Barro and Maizel, 2015)) and genes involved in lateral root emergence (*IDA*, *HAE* (Kumpf et al., 2013)), and *PIF3*. *HY5* and *HYH* itself were transcriptionally affected by GA_4_: *HYH* was downregulated in the shoot (Fig. S7a), while *HY5* was downregulated in the root (Fig. 8a), whereas HY5-GFP protein was enhanced (Fig. 07). This discrepancy between gene expression and protein response, may be related to different periods of GA exposure and the strong post-translational regulation of HY5 via COP1 (Lau and Deng, 2012).

**Figure 8.**
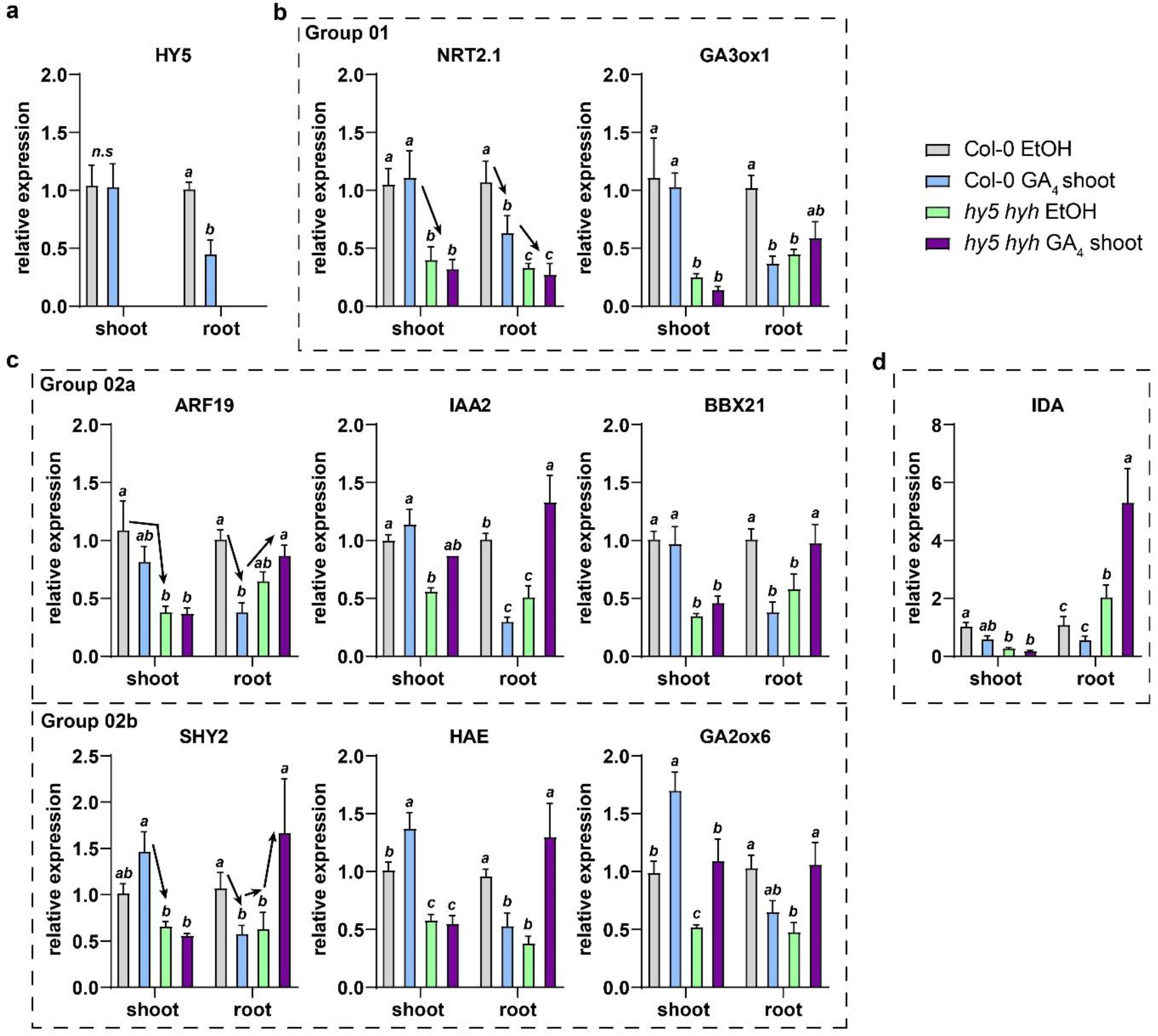
qPCR analysis shows repressive effect of GA4-shoot treatment on target gene expression in root, which is modulated by *hy5 hyh* mutation. qRT-PCR analysis of shoot and root samples of 5 day old seedlings (Col-0 and *hy5 hyh*), transferred to mock or GA4-shoot 24hrs before harvesting. (a) *HY5* expression values (not detectable in *hy5 hyh* mutant). (b) Group 01 genes, (c) Group 02a and 02b genes, (d) *IDA* expression values. Arrows show the changes in expression associated with that particular Group of genes. Expression values were analyzed according to the ΔΔ^CT^ method and normalized against the Col-0 mock sample for both shoot and root respectively. Four biological replicates per treatment, Error bars denote the standard error of the mean. Statistical significance was determined by 2-way ANOVA with a Newman-Keuls post-hoc test (p=0.05).

We were mainly interested in the effects of shoot-GA on gene expression in the root of wild type and the *hy5 hyh* mutant. Two genes that had a similar pattern of expression throughout the treatments were *GA3ox1* and *NRT2;1*; both showed reduced expression *hy5 hyh* compared to Col-0 and were downregulated by GA shoot treatment in the root tissues only of Col-0, but not *hy5 hyh* (Fig. 8b). As shown previously, the nitrate transporter gene *NRT2.1* is regulated via HY5 and is involved in the root WL+FR response (van Gelderen et al., 2021), while GA3ox1 is important in the later steps of GA biosynthesis (Hedden and Thomas, 2012).

A second, larger group was grouped on the basis of the root response. In this case GA-shoot caused a downregulation in the root, but in the *hy5 hyh* mutant roots these genes were then upregulated by GA (Fig. 8c, S7b, Group 2a). This group seems to be regulated by *GA*, but in this case the GA response in the root was modified by, but not completely dependent on, the *hy5 hyh* mutant. Four of these genes, (*SHY2, HAE, GA2ox6, GA2ox8*) were upregulated by GA in the shoot, while the rest (*ARF19, BBX21, IAA2, RGA*), were not (Fig. 8c, S7b, Group 2b). One gene, *IDA* (involved in lateral root emergence), was regulated particularly strongly via *HY5 HYH* since it was downregulated by GA in both shoot and root of Col-0, but its expression was enhanced in the *hy5 hyh* mutant and increased further in the GA-shoot treatment (Fig. 8d). Finally, a third group contained *PIF3* and *LAX3*, where the expression changes were relatively minor and not statistically significant (Fig. S7c). Overall, shoot GA_4_ treatment mainly had a repressive effect on the expression of the tested genes in the root. In most cases, the *hy5 hyh* mutation also resulted in a reduction in expression, meaning that in those cases HY5 would promote expression. However, in the *hy5 hyh* mutant the GA_4_-shoot effect was often opposite of that in wild type or was lost, indicating that *HY5* gates the GA response and highlighting the crosstalk between HY5 and GA.

To obtain more evidence for the role of HY5 in the downstream transcriptional response and at the same time the local effect of GA_4_, we performed an experiment with a *HY5* inducible line, driven by an estradiol-inducible cassette (Siligato et al., 2016), coupled to a lateral root primordium specific promoter (*pGATA23*) (De Rybel et al., 2010). This line was made in a wild type background to simulate the effect of a strong HY5 increase. At the same time, we added a paclobutrazol treatment in the same agarose solution used to elicit *pGATA23-XVE:HY5-YFP* expression in order to assess the effects of HY5 independent of putative interactions with endogenous GA.

reduced in expression by HY5 induction ((Fig. 9 b,d-e,g-h; Fig. S8), with the exception of *GATA22, NRT2.2* and *Ga3ox1* (Fig. S8). Interestingly, the PAC treatment did modulate some of the effects of the HY5 induction. *LAX3, SHY2, GA2ox6* and *RGA* were further repressed (Fig. 9d,e,g), while *GATA22* was further increased (Fig. S9) when GA was inhibited. In a similar experiment, but with the addition of a WL+FR treatment, we performed a ChIP-qPCR analysis to quantify the binding of HY5-YFP to the promoters of the genes tested for qPCR in Fig. 9 that also contained high affinity HY5 binding sites (Song et al., 2008). Since we used an inducible HY5-YFP line, we had three negative controls: the ’mock’ DMSO treatment (compared to estradiol), a general anti-igg antibody (compared to anti GFP/YFP) and a promoter without a HY5 binding site (*pFBP*). The beta estradiol induction of HY5 led to binding of HY5-YFP to the promoters of *HY5* itself (Fig. 9c). *HY5-YFP* induction led to binding of HY5 to the promoters of *NRT2.1* (Fig. 9c)*, LAX3* and *SHY2* (Fig. 9f), and *GA2ox6* and *HAE* (Fig. 9f). This coincided with a repression in expression of these genes (Fig. 9 a-b, d-e, g-h). Interestingly, WL+FR led to an increase in HY5-YFP binding for all genes concerned (Fig. 9c,f,i). For some genes PAC treatment led to a clear decrease in promoter binding of HY5-YFP (*GA2ox6, HAE* Fig. 9i), while for others PAC led to a small decrease (*HY5, NRT2.1, LAX3* and *SHY2* Fig. 9c,f). Thus, the specific HY5-YFP induction at the lateral root primordium development and initiation sites led to binding of HY5-YFP to the promoters of key genes for lateral root development and also to the promoter of a GA modifying enzyme. This was accompanied by a reduction in expression of these genes. PAC treatment had a mild effect on HY5 promoter binding and expression effects, while WL+FR treatment led to an increase in promoter binding. In many of the same genes, shoot-GA_4_ treatment led to a repression in expression, in accordance with the positive effect of GA on HY5 protein amounts, while the effect of shoot-GA_4_ effect on gene expression was reversed in the *hy5 hyh* mutant.

**Figure 9.**
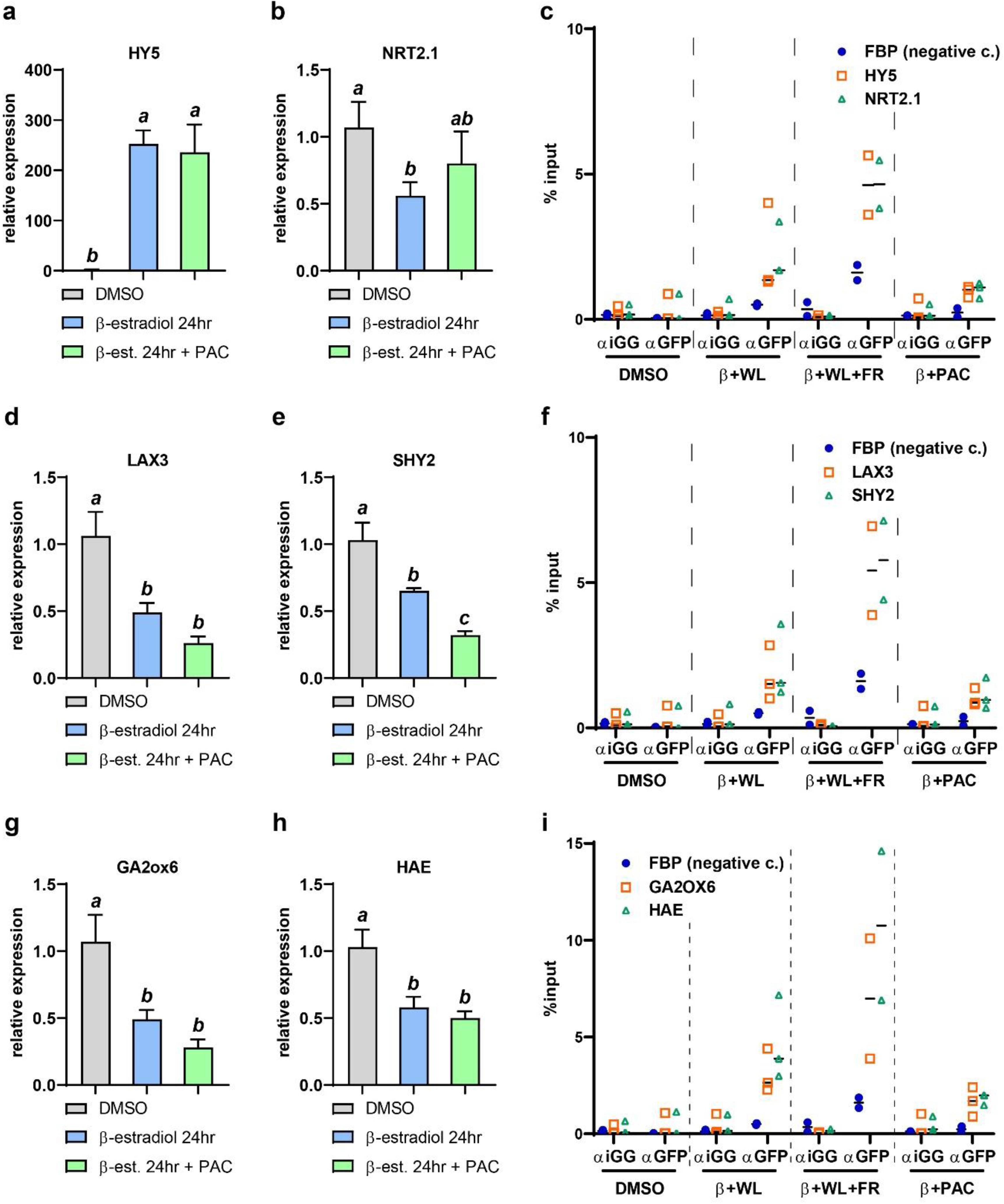
A qCPR and ChIP-qPCR approach shows repressive effect of HY5 on expression of lateral root development and GA-related genes. (a-b, d-e, g-h) qRT-PCR analysis of root samples of 5 day old seedlings (Col-0 *pGATA23-XVE:HIS-HY5-YFP*), induced on the root by beta-estradiol, mock induced and treated with paclobutrazol (PAC) 24hrs before harvesting. Expression values were analyzed according to the ΔΔ^CT^ method and normalized against the Col-0 mock sample for both shoot and root respectively. Four biological replicates per treatment, Error bars denote the standard error of the mean. Statistical significance was determined by 1-way ANOVA with a Newman-Keuls post-hoc test (p=0.05). (c,f,i) Chromatin Immuno Precipitation (ChIP) – qRT-PCR experiment on the same Col-0 *pGATA23-XVE:HIS-HY5-YFP* line, with 5 day old seedlings, treated 24 hours earlier with mock, beta-estradiol with additional WL+FR or PAC treatment. Roots were sampled and the isolated and sheared chromatin was incubated with either a control anti-iGG antibody, or an anti-GFP antibody, probing for HY5-YFP. qRT-PCR values were calculated to a percentage of the input-DNA before immunoprecipitation.

## Discussion

We have shown previously that above-ground competitive responses can lead to a reduction in root growth, by a reduction in lateral root emergence regulated through HY5 (van Gelderen et al., 2018) and that this response depends upon the nutrient status of the plant (van Gelderen et al., 2021). Importantly, in these studies only the shoot is in an environment with a low R:FR ratio, since the roots are shielded from the far-red light via a cover around the plate and barrier in the plate (van Gelderen et al., 2018). Now we show that these responses are modulated via Gibberellin. Whole-plate GA application has shown that lower GA_4_ levels modulate the root response to far-red light application on the shoot (WL+FR) and that higher GA_4_ levels inhibit root growth. When GA biosynthesis is blocked, via PAC application, we observed a reduction in root growth and a loss of the WL+FR phenotype. The reason that both GA_4_ addition and the block of GA biosynthesis can give similar phenotypes at certain concentrations, may lie in the fact that root growth can be repressed by ectopic expression of DELLAs (Heo et al., 2011b; Zhang et al., 2011), mimicking PAC application, but that normal DELLA signaling is also necessary to allow root growth (Ubeda-Tomás et al., 2009, 2008), which is important when applying physiologically relevant levels of GA.

With the use of the GPS1 GA-FRET sensor (Rizza et al., 2021, 2017), we were able to show that GA is increased in the lateral root primordia and in the elongation zone of the root. Shoot-application of GA_4_ and the GA-biosynthesis inhibitor uniconazole has previously been shown to reduce root growth (Bidadi et al., 2010). In the shoot, gibberellin biosynthesis is upregulated and gibberellins accumulate in low R:FR (Djakovic-Petrovic et al., 2007; Kohnen et al., 2016). Gibberellins are essential regulators in hypocotyl growth (Rizza et al., 2017) and cooperate with auxin in the low R:FR response of adult plant petioles (Küpers et al., 2023). GA_3_ and GA_4_ can be transported from shoot to root through the phloem (Shani et al., 2013). In our experiments, we have shown that shoot application of GA_4_ leads to a reduction in root growth, but more importantly, also to a loss of the WL+FR effect. Shoot-applied GA_4_ led to a clear response of the GPS1 sensor in the root: in the vasculature, lateral root primordium, and the elongation zone. Thus, our data together indicate that shoot-root transport of GA plays an important role in the root response to WL+FR. The question is then how shoot-derived GA_4_ affects the development of the root. Gibberellin signaling acts through the proteasomal degradation of the DELLA transcriptional regulators. The *della pentuple* mutant did not have a WL+FR root phenotype, and did not respond to GA application, thus supporting the notion that the GA_4_ response is mediated by DELLAs. DELLAs frequently act by directly binding to transcription factors whose activity is then inhibited (Daviere and Achard, 2016) and they can affect main root growth through the interaction of RGA with SCARECROW-LIKE3 (Heo et al., 2011a; Zhang et al., 2011). GA_3_ (thus presumably GA_4_ as well) can enhance the responsiveness of the root to IAA (Li et al., 2015). In our experiments, GA_4_-shoot treatment also led to a decrease in detected auxin (R2D2), auxin signaling (dr5v2) and a repression of auxin-related gene expression (qPCR). GA can also promote auxin signaling by relieving DELLA repression of ARF6 and ARF8 (Oh et al., 2014), and to understand the effects of GA on root auxin response gene expression, future studies could investigate if DELLA also interacts with ARFs that are more involved in the regulation of root auxin levels and root development, such as ARF7 and ARF19.

The *hy5 hyh* double mutant, which showed not root response to WL+FR treatment, was also less responsive to GA_4_ treatment, suggesting that GA_4_ response in roots involves HY5. GA_4_-shoot treatment led to an increase of HY5 in the root and HY5 is known to be able to repress auxin signaling in the root (Cluis et al., 2004; Sibout et al., 2006; van Gelderen et al., 2018), which was confirmed by the qPCR and ChIP-qPCR data after HY5 induction. Furthermore, the qPCR results showed that in the *hy5 hyh* mutant, the effect of GA_4_ application was modified: In *hy5 hyh* roots, GA_4_-shoot treatment led to an increase in auxin-related expression, indicating that HY5 normally represses this GA response. Together these results paint a picture whereby low R:FR detected by the shoot, leads to increased gibberellin transport to the root, either or not in parallel with putative HY5 translocation. In the root gibberellin stimulates HY5, which leads to repression of auxin signaling and thus of (lateral) root growth (Fig. 10).

**Figure 10.**
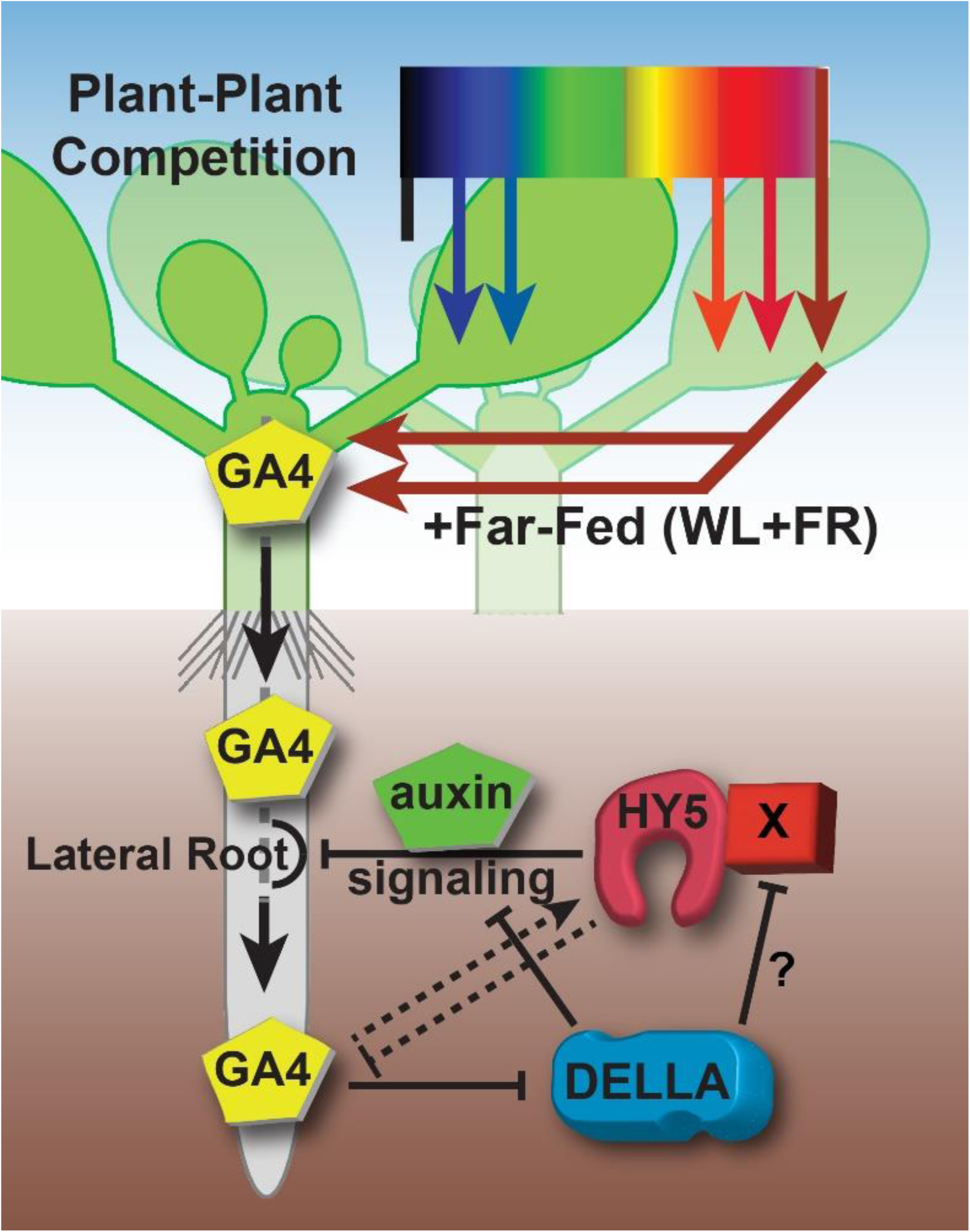
The role of Gibberellins (GA4) and HY5 in modulating lateral root growth during above-ground plant-plant competition. The proximity of neighboring plants leads to a relative increase in the amount of far-red light, simulated in our experiments with the addition of far-red light to the growth light (WL+FR). This leads to an increase in GA, which can travel through the phloem towards the root tissues. There, GA leads to DELLA degradation, which indirectly leads to an increase in HY5, possibly via the de-repression of HY5 cofactors or an as-of-yet unidentified mechanism. Alternatively, DELLA could repress auxin signaling directly by binding to ARF transcription factors. HY5 can repress auxin signaling and thus a brake on lateral and main root development. However, HY5 can also repress GA biosynthesis.

An open question is then how gibberellin leads to a stimulation of HY5 protein levels. Until now, there has been no evidence that RGA or any other DELLA interacts with HY5. RGA can bind to and inhibit PIF3 and BZR1, key regulators of light and hormone signaling (Bai et al., 2012; De Lucas et al., 2008; Gallego-Bartolome et al., 2012) that could be interesting collaborative factors for HY5 action, since PIF3 and BZR1 can interact with HY5 and act as a cofactor of HY5 (Chen et al., 2013; Li and He, 2016). Future studies could investigate the interactions that occur between HY5 and other transcriptional regulators in the root, to really pinpoint the mechanism of regulation.

## Materials and Methods

### Plant material and growth conditions

*Arabidopsis thaliana* plants were grown in a controlled environment growth chamber with a 16-h-light/8-h-dark cycle, temperature of 20°C and a light level of PAR ∼140 µmol/m^2^/s. Knockout lines used were: *hy5 hyh* (Zhang et al., 2017)*, dellap pentuple (gai-t6 rga-t2 rgl1-1 rgl2-1 rgl3-4), pif1 pif3 and pif1 pif3 pif4 pif5 pif7* (Leivar et al., 2020)*, ga1-3* (Michaels and Amasino, 1999). Fluorescent marker lines and sensors used were: *NLS-GPS1* and *NLS-GPS1-NR* (Rizza et al., 2017)*, hy5-2 pHY5:HY5-GFP* (Zhang et al., 2017)*, pRGA:GFP-RGA* (Silverstone et al., 2001)*. C3PO* (Küpers et al., 2023). Col-0 *pGATA23-XVE:HIS-HY5-YFP* was created by stably transforming Col-0 with a tissue specific vector system (Siligato et al., 2016), harbouring a newly cloned *HY5* coding sequence (see molecular cloning).

### Square plate growth and root phenotyping

For root phenotyping seeds were surface sterilized using a 1 hour treatment with chlorine gas. Seeds were sown on ½ MS 0.1% MES, pH 5.8, 0.8 % plant agar plates, at 25 per plate (12,5x12,5x1,75 cm) by scattering them on a plate and then the plates were put in the dark at 4°C for six days for seed stratification. Plates were put in the growth chamber (16/8 light/dark photoperiod, PAR = 140 µmol m^-2^ s^-1^, 21 °C) shortly after subjective dawn and allowed to germinate for 24 hours. Then the germinated seeds were transferred on a new plate, containing either the treatment chemical or a mock treatment, 25 seeds on a line, at 9 cm height. Just below the seeds the D-root (Silva-Navas et al., 2015) insert was placed and the root part of the plate was covered by a black cover (see (van Gelderen et al., 2018)). Plates were then returned to the growth chamber after which either white light treatment continued or the WL+FR treatment started. For the WL+FR treatment plates were placed 20 cm in front of a row of FR LEDs (Phillips GreenPower LED research module far red, 24Vdc/10W) to achieve a R:FR ratio of 0.1 in the shoot part, which was measured inside the plate using a small, flexible R:FR light meter (Skye Spectrosense2). After four days of growth, seedlings were transferred in the afternoon to a new plate (5-6 seedlings per plate) to ensure homogeneous growth and prevent intermingling of root systems. Square Petri dishes were scanned at 600 or 1200dpi using an EPSON V850 photonegative scanner. Experiments were analyzed using Smartroot (Lobet et al., 2011). The scans were converted from a color image to an 8bit grayscale image and contrast was enhanced to provide a better input for Smartroot. Hypocotyl length was analysed either manually with a precision calliper after scanning, or by analysing the scanned images with imageJ.

### DIC microscopy and lateral root primordia analysis

Seedlings used for lateral root primordia analysis were fixed and cleared according to a previously published protocol (Malamy and Benfey, 1997). The seedlings were mounted in 50% glycerol and analyzed using a Zeiss Axioskop2 DIC microscope (40x Plan-NEOFLUAR DIC objective) with a Lumenera Infinity 1 camera.

### Confocal microscopy and analysis

For confocal microscopy seeds were sown at 16 per plate and were transferred at day 4 to GA_4_ or PAC containing medium. Seedlings were fixed with 4% paraformaldehyde and cleared and stained according to a modified clearsee protocol (Ursache et al., 2018). Confocal microscopy was performed with either a Zeiss Observer Z1 LSM7 confocal imaging system, with 405, 488, 514 and 563 nm excitation lasers, and fixed bandpass filters for CFP, YFP and RFP, or a Zeiss LSM 880 Airyscan system using 453, 488, 514 and 563 lasers with electronically adjustable bandpass filters. Confocal z-stack images were made using a 40x NA1.2 oil immersion objective (both microscopes). Within experiments, pinhole, detector gain, laser power and detector offset were the same. All images were analyzed using ICY http://icy.bioimageanalysis.org/), using the HK-means plugin to select individual nucleus regions.

### RNA extraction and qRT-PCR expression analysis

For gene expression analyses, plants were sown at 16 seeds in a row and grown in the conditions mentioned above for 5 days. Between 15 to 19 seedlings were harvested per sample (from two plates) and only root tissues were used for RNA extraction. Four biological replicates were taken per treatment/genotype condition. The Qiagen plant RNeasy kit was used for RNA extraction. First-strand cDNA was made using the Thermo Scientific RevertAid H Minus Reverse Transcriptase, RiboLock RNase inhibitor, and Invitrogen random hexamer primers. RNA input into the cDNA reaction was kept equal within experiments, preferably at 1000 ng. Primers were designed preferably across introns and for 100- to 150-bp fragments with an annealing temperature of 60°C with primer3plus (http://www.bioinformatics.nl/cgi-bin/primer3plus/primer3plus.cgi). Primers were tested for efficiency using generic Col-0 cDNA at a concentration range of 2.5→40 ng of cDNA per 5 mL reaction. qPCR reagents used were Bio-Rad SYBR-Green Mastermix on 384-well plates in a Life Technologies ViiA7 real-time PCR system. All CT values were normalized against two validated housekeeping genes: *ADENINE PHOSPHORIBOSYL TRANSFERASE1 (APT1) and PROTEIN PHOSPHATASE 2A SUBUNIT A3 (PP2AA3)*. The ΔΔ^CT^ method was used to calculate relative expression values (Livak and Schmittgen, 2001). Primer sequences are provided in the supplementary table.

### Chromatin immunoprecipitation

For the Chromatin Immuno Precipitation (ChIP) experiment we used a modified version of a histone modification ChIP-seq protocol (Antunez-Sanchez et al., 2020; Ramirez-Prado et al., 2021). We used ∼150-200 seedling roots per sample, and performed the chromatin isolation in 2ml eppendorf tubes, rather than falcon tubes. Shearing was done with a Diagenode Bioruptor, for a 20-30 minute length of 30 second bursts, followed by 30 seconds of rest. The immunoprecipitation was performed in PCR strips, using Pierce protein A/G magnetic beads, Roche anti GFP monoclonal antibody and Roche rabbit anti-igg antibody as control. Isolated DNA fragments were measured with a qBit DNA analyzer. For the qRT-PCR reactions, 20-50 picogram of DNA was used per reaction, with the same reagents as described above.

### Molecular cloning and plant transformation

We made a transgenic, estradiol-inducible, tissue-specific HY5 line (Col-0 *pGATA23-XVE:HIS-HY5-YFP*). We cloned the *pGATA23* promoter into an empty multisite gateway-compatible entry vector (p1R4-pGATA-XVE), by first PCR-amplifying a *pGATA23* fragment and with BshTI and XhoI restriction sites on the 5’ ends of the primers. We ligated this fragment into a pJet1.2 cloning vector. From this vector we excised and ligated the *pGATA23* fragment into p1R4-ML-XVE (Siligato et al., 2016). We created a pDONR207-HIS-HY5 entry vector by PCR amplifying from cDNA a HY5 fragment with flanking gateway sites and performing a BP reaction. We then performed a multisite gateway reaction using the entry vector with the p1R4-pGATA-XVE, pDONR207-HIS-HY5 and p2R3 YFP 3AT entry vectors in a pGII R3R4 entry vector (Siligato et al., 2016) (Addgene). Primers are shown in the supplementary table. This entry vector was then transformed to dh5a *E.coli,* and subsequently prepped and transformed into *AGL1 psoup A. tumefaciens.* The transformation of Arabidopsis flowering plants was done via a published floral dip method (Davis et al., 2009). Transformed T1 seeds were selected with hygromycin and propagated. ∼20 T2 lines were selected based on mendelian segregation to select for single inserts and also tested with epifluorescence microscopy for leaky expression and fast and correct induction. ∼5 T3 lines were selected for homozygosity with hygromycin and again tested for proper, fast non-leaky induction.

### Data processing and statistics

Data from Smartroot and Icy was processed and analysed with R with a custom-made script. Two-way ANOVA testing with post-hoc tukey tests was done with R. Graphs were made using Graphpad Prism and One-way ANOVA and t-test were also done with Graphpad Prism. Boxplot graphs show min/max excluding outliers, with outliers identified by a tukey test as individual data points. Figures were composed using Adobe Illustrator.

### Author Contributions

K.v.G. and R.P. designed the experiments and wrote the manuscript. K.v.G., K.v.d.V., C.K., J.H., O.P., A.K. and J.H. performed and analyzed the experiments. T.A performed experiments and provided technical assistance.

## Acknowledgements

We thank Richard Paalman, Roxanne Lock and Alicia Koppenol for experimental optimisations. We thank Leonardo Jo for his help with setting up ChIP-qPCR experiments and the Utrecht Scientific Instrument Workshop for their assistance with manufacturing custom plates. We thank Alexander Jones for sharing GPS1 lines and their guidance on GPS1 imaging, and Lanlan Zheng for sharing a collection of *HY5* lines.

## Funding Information

This research was funded by the Netherlands Organisation for Scientific Research, open competition grant 823.01.013 and Vici grant 865.17.002 to R.P. CK was funded by a scholarship of government sponsorship for overseas study, Taiwan, admission number 0991167-2-UK-004. KvG is currently funded by the Emmy Noether program of the DFG #GE 3355/1-1.

## Notes

### Competing Interest Statement

The authors have declared no competing interest.

