## Supplemental Figures for "Gibberellin transport affects (lateral) root growth through HY5 during Far-Red light enrichment"

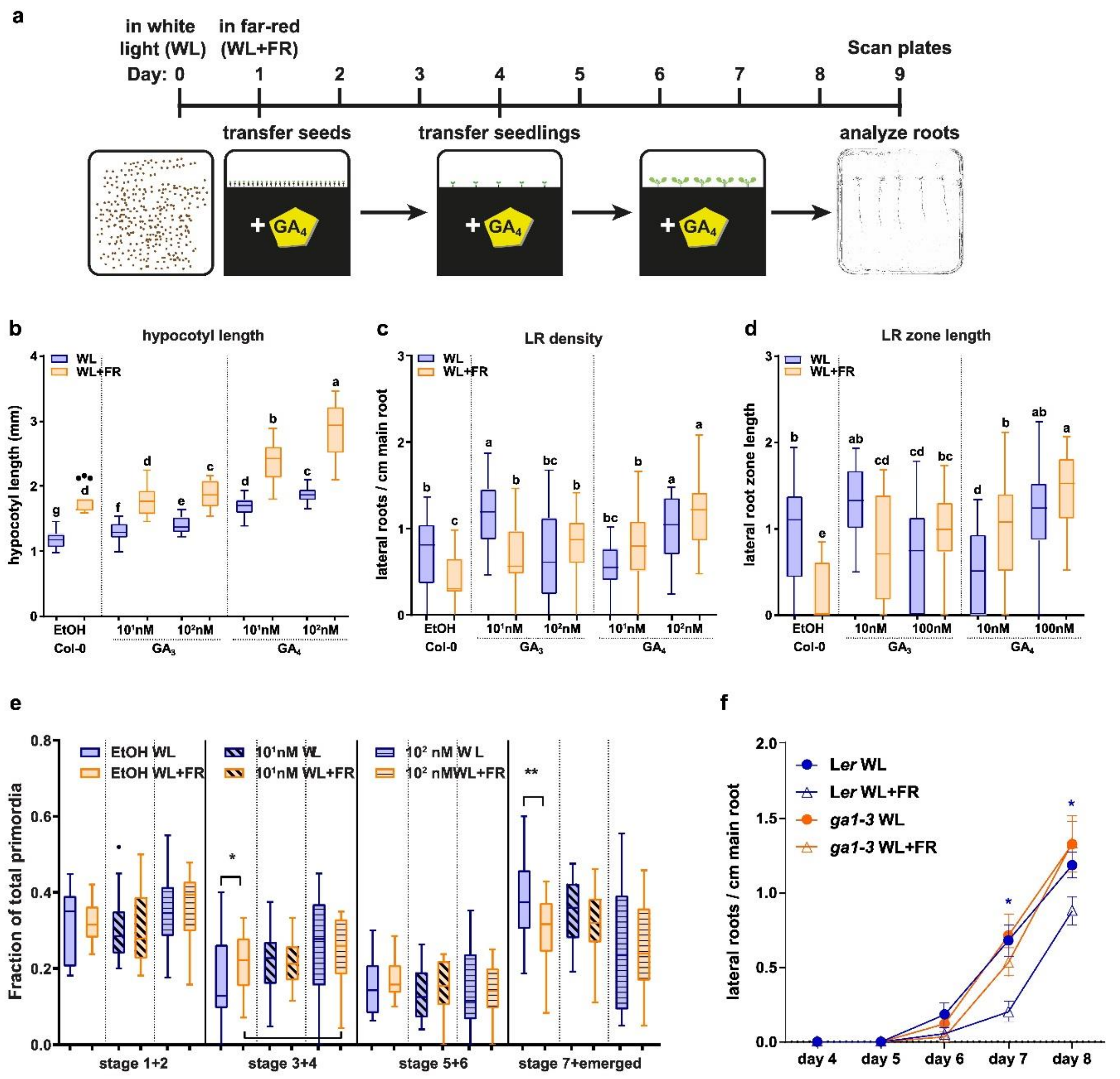

**Supplemental Figure 1 | Gibberellins are involved in FRshoot-induced changes in root development.** (a) Growth scheme for phenotyping experiments of Figures 1,2,6,10 and S1, S4, S6. (b-d) Seedlings were grown for 9 days according to the schedule on (a). Scans of 8d old seedlings were analyzed on (b) hypocotyl length; (c) lateral root density; (d) lateral root zone length. (e) A similar experiment to Figure 1 was performed with GA4 addition, however seedlings were fixed afterwards and analyzed with DIC microscopy on lateral root primordia stages. (f) Lateral root density of experiment performed similarly to the schedule in (a), with the addition that seeds were surface sterilized with ethanol and bleach and then taken up in 0.1% agarose, which for the *ga1-3* mutant contained $10^2$ nM of GA4, in order to start germination of this otherwise non-germinating line. Means were statistically significant based on a 2-way ANOVA; letters denote significant difference between treatments based on a post-hoc tukey test ($p<0.05$).

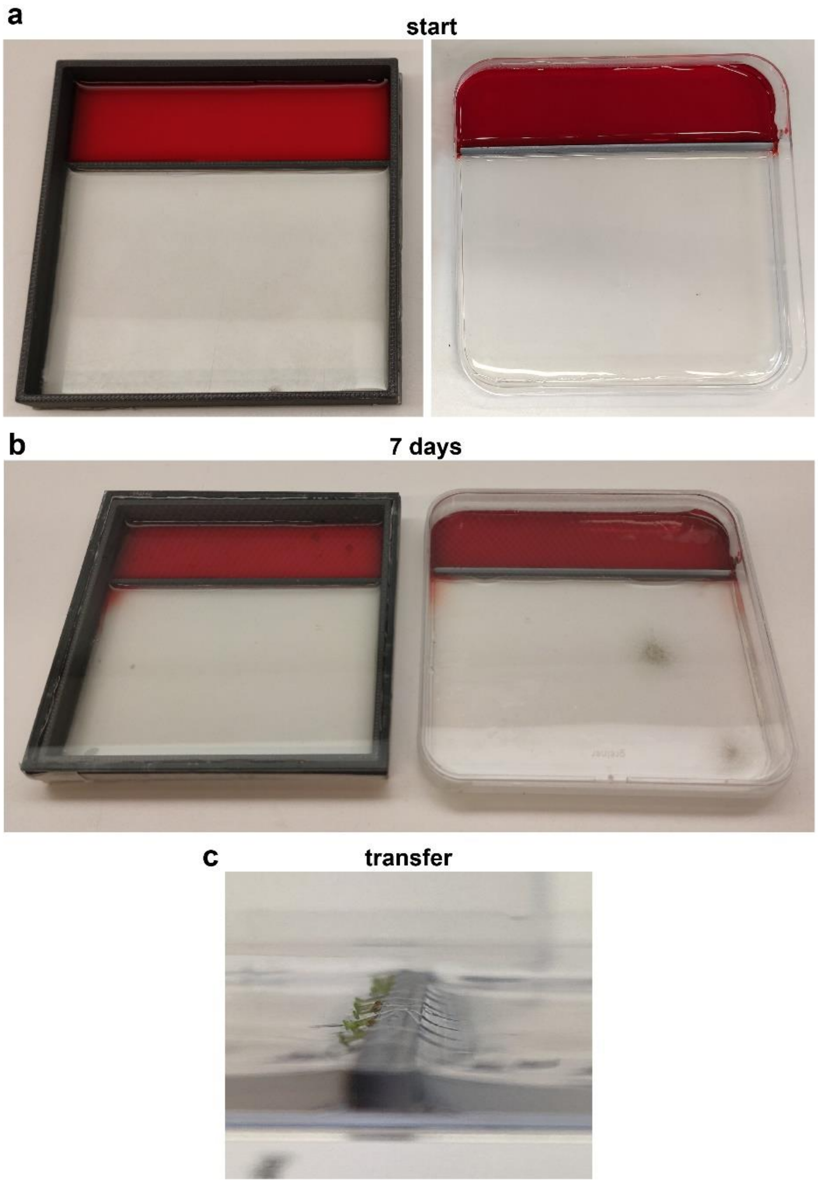


**Supplemental Figure 2 | Compartmentalized plates have minimal diffusion after seven days of vertical orientation and seedlings are carefully positioned across the border.** Two types of two-compartment plates were constructed and tested. On the left a 3D printed plastic frame was glued to a glass bottom, with an accompanying 3D printed lid, also glued to a glass plate. The second plate (right), was made by gluing with epoxy-resin a plastic insert of 3mm into an off-the-shelf Greiner 12 cm square plate. (b) Two agar layers were poured into the plates, a top one containing 0.1% direct red 23, and a bottom one, without any addition. After 7 days of vertical placement, which is two days longer than the running time of the experiment, only minimal leakage of the dye on the sides of the plate had occurred, demonstrating that the two layers can stay sufficiently separated. (c) Close-up image of four-day old seedlings transferred onto the two-compartment plate, with the hypocotyl and shoot part touching the top layer, and the root part touching the bottom layer.

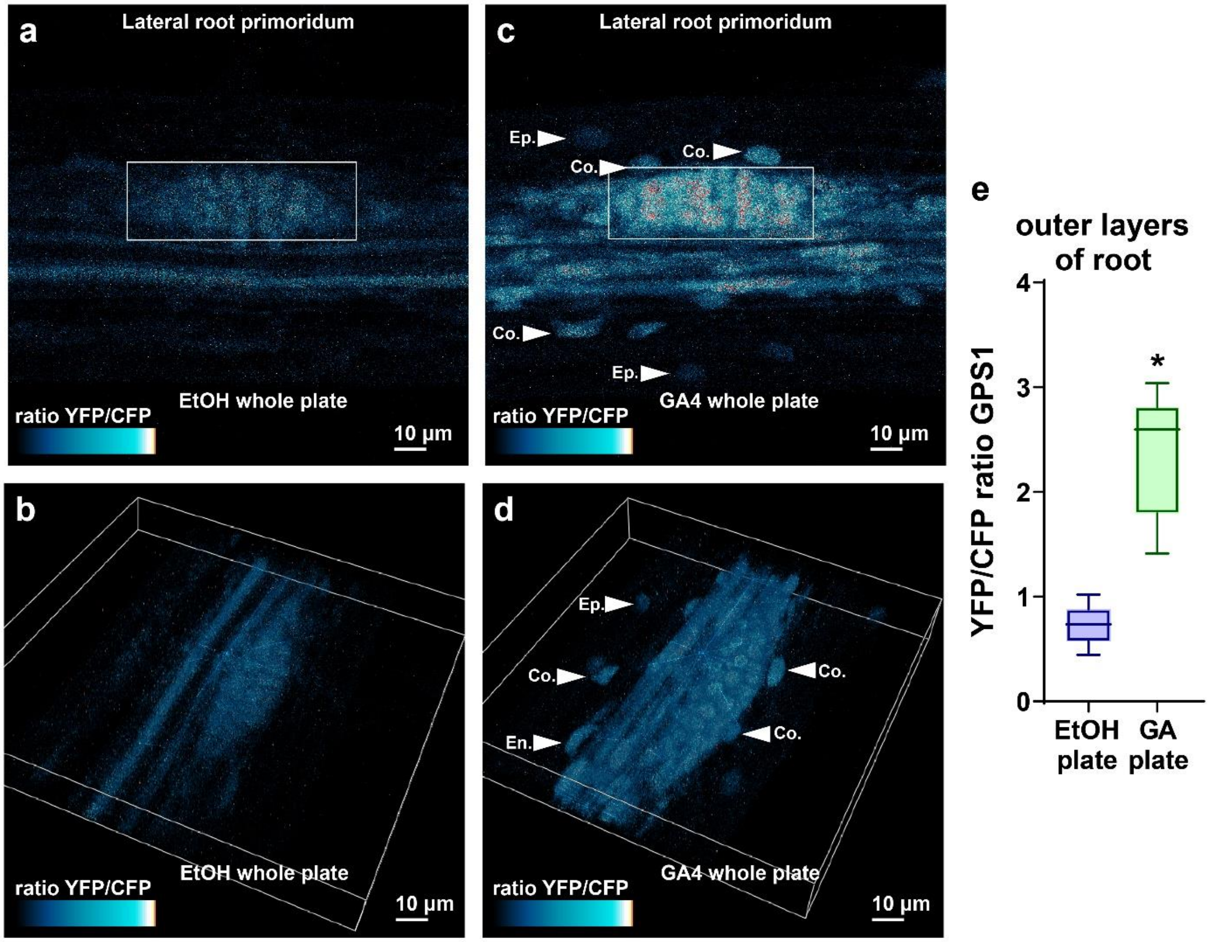


**Supplemental Figure 3 | GPS1 ratios in the outer root layers (cortex, epidermis) increase strongly with a whole plate GA4 treatment.** (a-d) Representative images of stage 4 lateral root primordia of six day old seedlings from the GPS1 line treated with mock or GA4. Shown is the ratio between YFP and CFP emission of GPS1 (a,c) 2D projection, (b,d) 3D projection, White box emphasizes the LRP, and nuclei of the epidermis (Ep.), cortex (Co.) and endodermis (En.) layers have been highlighted .white arrow denotes cortex and epidermis cell nuclei with an increase in GPS1 ratio. (e) Quantification of images from the whole experiment. *significant difference based on student's t-test.

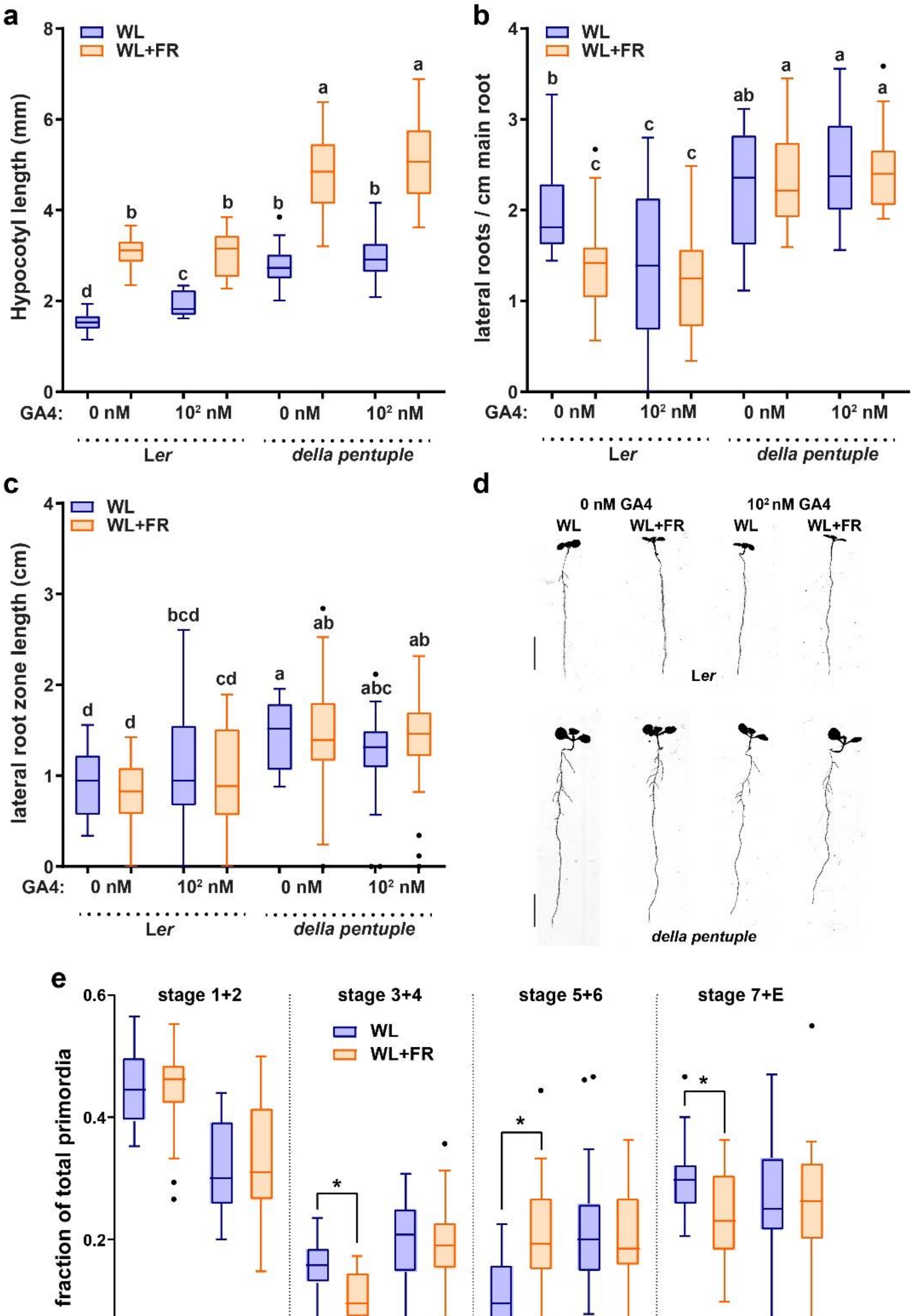


**Figure 4 | The *della pentuple* mutant has no lateral root development reduction in WL+FR and does not respond to a GA4 treatment.** Seedlings of L*er* and *della pentuple* were grown for 9 days according to the schedule on Fig. S1a. Scans of 9d old seedlings treated with EtOH 1:10000 in the medium or $10^2$ nM GA4 and analyzed on (a) hypocotyl length; (b) lateral root density; (c) lateral root zone length. (d) Representative seedling images of the experiment in (a-c). (g-l) Similar setup as in (a-f), but then with 10 μM GA4. scale bar = 1 cm. Means were statistically significant based on a 2-way ANOVA; letters denote significant difference between treatments based on a post-hoc tukey test ($p<0.05$).

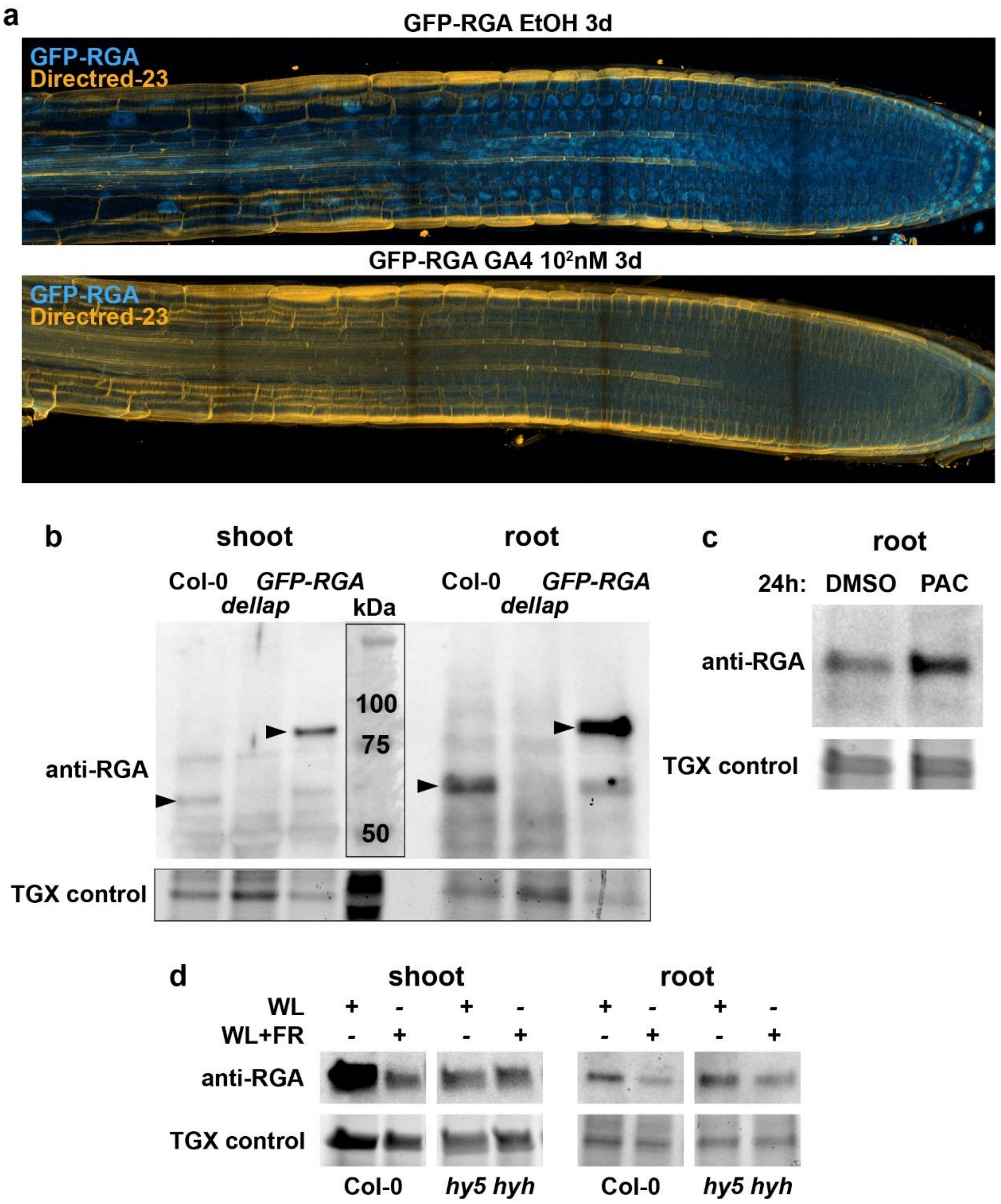


**Supplemental Figure 5 | Additional experiments show that *della pentuple* mutant does not react to GA and has a reduced root Far-Red response.** (a) Microscopy images of 7 day old seedling roots of *rga pRGA:GFP-RGA*, treated with mock EtOH or with $10^2$ nM of GA4. (b) Western blot experiment using 5 day old seedling shoots and roots of Col-0, *della pentuple* and of *rga pRGA:GFP-RGA,* demonstrating the specificity of the RGA antibody. (c) Western blot experiment using 5 day old seedling roots of Col-0 treated for 24 hours with mock (DMSO) or 1 µM paclobutrazol. (d) Western blot experiment using 5 day old seedling shoots and roots of Col-0 and *hy5 hyh,* grown either in WL, or in WL+FR (for 24 hours).

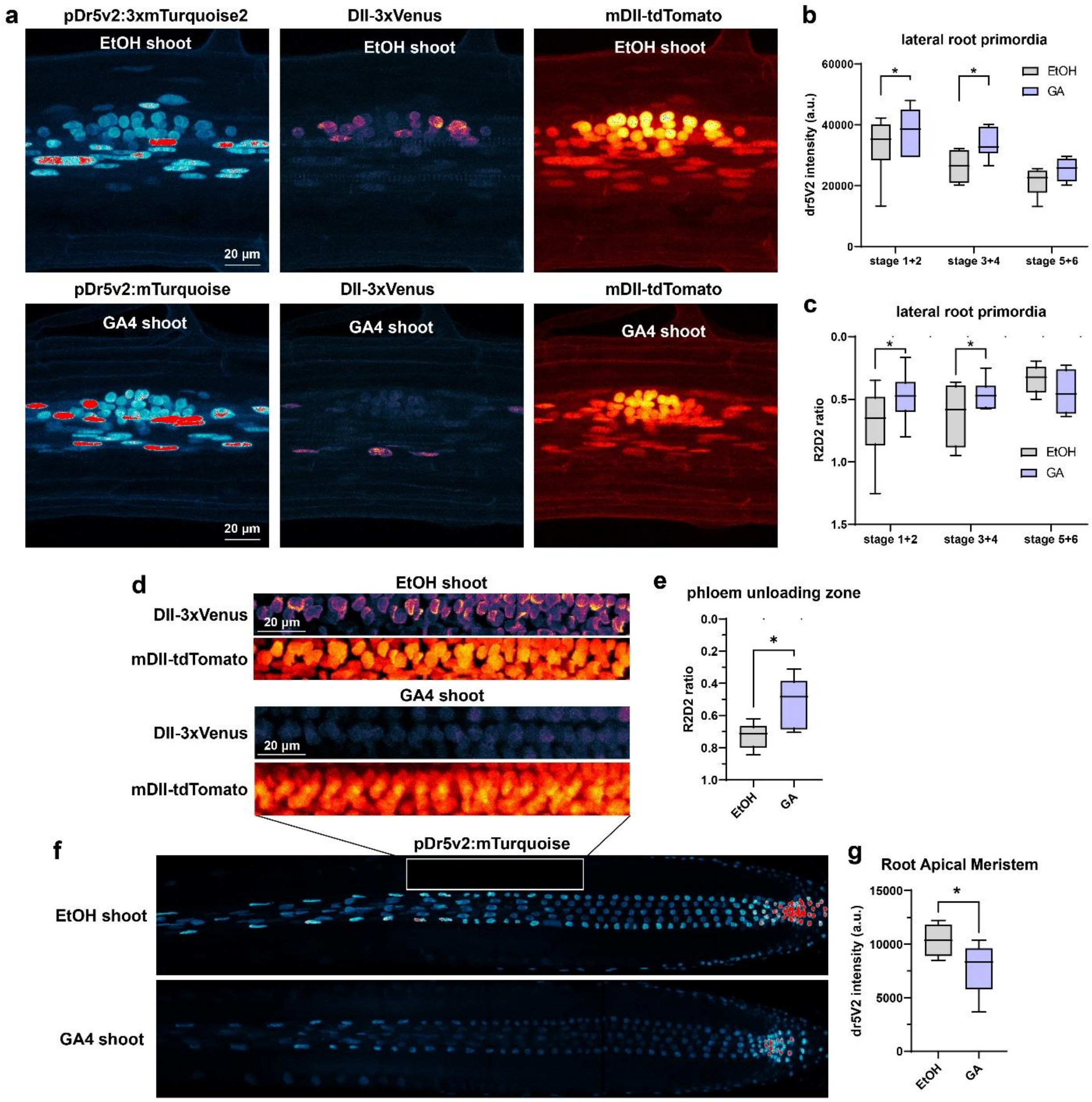


**Supplemental Figure 6 | GA4 shoot treatment leads to changes in auxin levels and signaling in root tissues.** Confocal microscopy experiment, comparable to Fig. 5, with the C3PO line, containing the R2D2 auxin level sensor and the Dr5v2 auxin signaling sensor. (a,b) C3PO line was either treated for two days with shoot GA4, or with mock EtOH, representative images of a stage 4 lateral root primordium (a) and quantifications of the dataset (b,c). (d) Representative images of R2D2 in the phloem unloading zone (for reference, see white box in (f)), with corresponding quantifications (e). (f) Representative images of *pDR5v2:mTurquoise* in the main root tip, white box denotes the phloem unloading zone, with quantification of dataset in (g) * significant difference based on student's t-test ($p<0.05$).

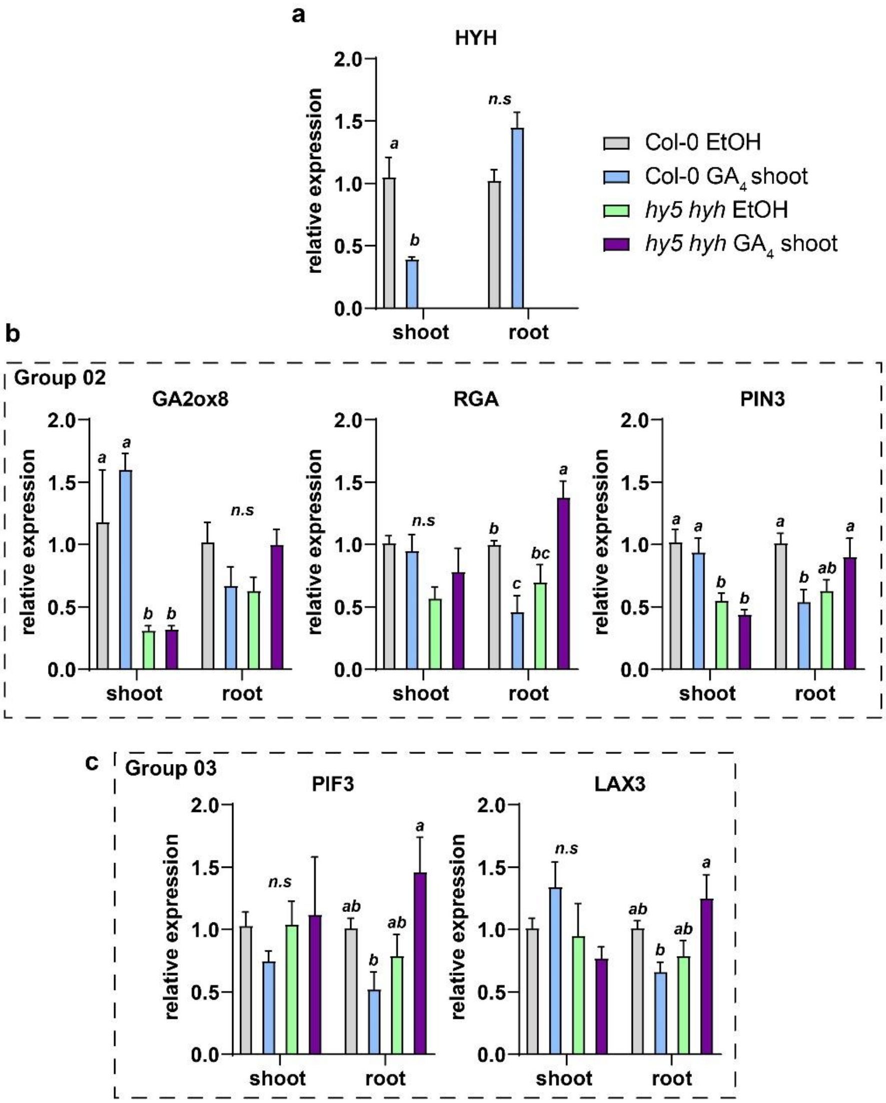


**Supplemental Figure 7 | qPCR experiment with shoot GA4 treatment and *hy5 hyh* mutant, supplementary to Figure 8.** qRT-PCR analysis of shoot and root samples of 5 day old seedlings (Col-0 and *hy5 hyh*), transferred to mock or GA4-shoot 24hrs before harvesting. (a) *HYH* expression values (not detectable in *hy5 hyh* mutant). (b) Additional cluster 02 genes, (c) Cluster 03 genes. Expression values were analyzed according to the $\Delta\Delta^{CT}$ method and normalized against the Col-0 mock sample for both shoot and root respectively. Four biological replicates per treatment, Error bars denote the standard error of the mean. Statistical significance was determined by 2-way ANOVA with a Newman-Keuls post-hoc test (p=0.05).

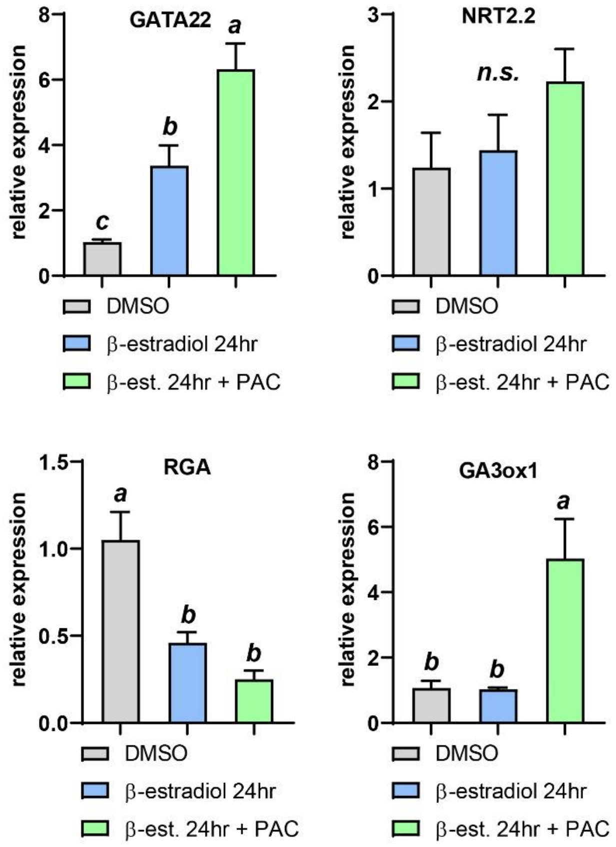


**Supplemental Figure 8: Additional qPCR data to Figure 9** qRT-PCR analysis of root samples of 5 day old seedlings (Col-0 *pGATA23-XVE:HIS-HY5-YFP*), induced on the root by beta-estradiol, mock induced and treated with paclobutrazol (PAC) 24hrs before harvesting. Expression values were analyzed according to the $\Delta\Delta^{CT}$ method and normalized against the Col-0 mock sample for both shoot and root respectively. Four biological replicates per treatment, Error bars denote the standard error of the mean. Statistical significance was determined by 1-way ANOVA with a Newman-Keuls post-hoc test (p=0.05).
